# Specification and survival of post-metamorphic branchiomeric neurons in the hindbrain of a non-vertebrate chordate

**DOI:** 10.1101/2023.06.16.545305

**Authors:** Eduardo D. Gigante, Katarzyna M. Piekarz, Alexandra Gurgis, Leslie Cohen, Florian Razy-Krajka, Sydney Popsuj, Hussan S. Ali, Shruthi Mohana Sundaram, Alberto Stolfi

## Abstract

Tunicates are the sister group to the vertebrates, yet most species have a life cycle split between swimming larva and sedentary adult phases. During metamorphosis, larval neurons are largely replaced by adult-specific ones. Yet the regulatory mechanisms underlying this neural replacement remain largely unknown. Using tissue-specific CRISPR/Cas9-mediated mutagenesis in the tunicate *Ciona*, we show that orthologs of conserved hindbrain and branchiomeric neuron regulatory factors Pax2/5/8 and Phox2 are required to specify the “Neck”, a compartment of cells set aside in the larva to give rise to cranial motor neuron-like neurons in the adult. Using bulk and single-cell RNAseq analyses, we also characterize the transcriptome of the Neck downstream of Pax2/5/8. Surprisingly, we find that Neck-derived adult ciliomotor neurons begin to differentiate in the larva, contrary to the long-held assumption that the adult nervous system is formed only after settlement and the death of larval neurons during metamorphosis. Finally, we show that manipulating FGF signaling during the larval phase alters the patterning of the Neck and its derivatives. Suppression of FGF converts Neck cells into larval neurons that fail to survive metamorphosis, while prolonged FGF signaling promotes an adult neural stem cell-like fate instead.

## Introduction

The simple embryos of the non-vertebrate chordate *Ciona* and related tunicates comprise a highly tractable system in which to study the regulation of cellular processes in development (Farley et al. 2015; Bernadskaya and Christiaen 2016; Cao et al. 2019; Wang et al. 2019; Guignard et al. 2020). Their classification as tunicates, the sister group to the vertebrates (Delsuc et al. 2006), means they share with vertebrates many chordate-specific gene families, cell types, organs, and anatomical structures (Christiaen et al. 2002; Dufour et al. 2006; Razy-Krajka et al. 2014; Abitua et al. 2015; Stolfi et al. 2015; Di Gregorio 2020; Fodor et al. 2021; Lemaire et al. 2021; Papadogiannis et al. 2022). The simplicity of their embryos overshadows the fact that the larva is but one part of a biphasic life cycle alternating between a free-swimming larval phase and a sessile adult phase (**Figure 1A**). During metamorphosis, larval structures degenerate and are replaced by adult tissues and organs (Nakayama-Ishimura et al. 2009; Sasakura and Hozumi 2018). Although the invariant cell lineages and gene regulatory networks specifying many of the embryonic cell types of *Ciona* have been investigated in detail (Imai et al. 2006; Cao et al. 2019), metamorphosis and adult development are still poorly understood. We know that various undifferentiated progenitor cells set aside in the larva as discrete compartments can give rise to adult structures (Dufour et al. 2006; Horie et al. 2011; Razy-Krajka et al. 2014; Sasakura and Hozumi 2018). These adult progenitor cells are patterned and specified in an invariant manner alongside differentiated larval cells, but only fully differentiate after the larva finds a location to settle and undergo metamorphosis. This transition from invariant (stereotyped) to variable (plastic) development is unique among the chordates, and thus of potential interest for revealing previously unknown mechanisms for precise spatiotemporal regulation of cellular quiescence, survival, proliferation, and differentiation.

**Figure 1.**
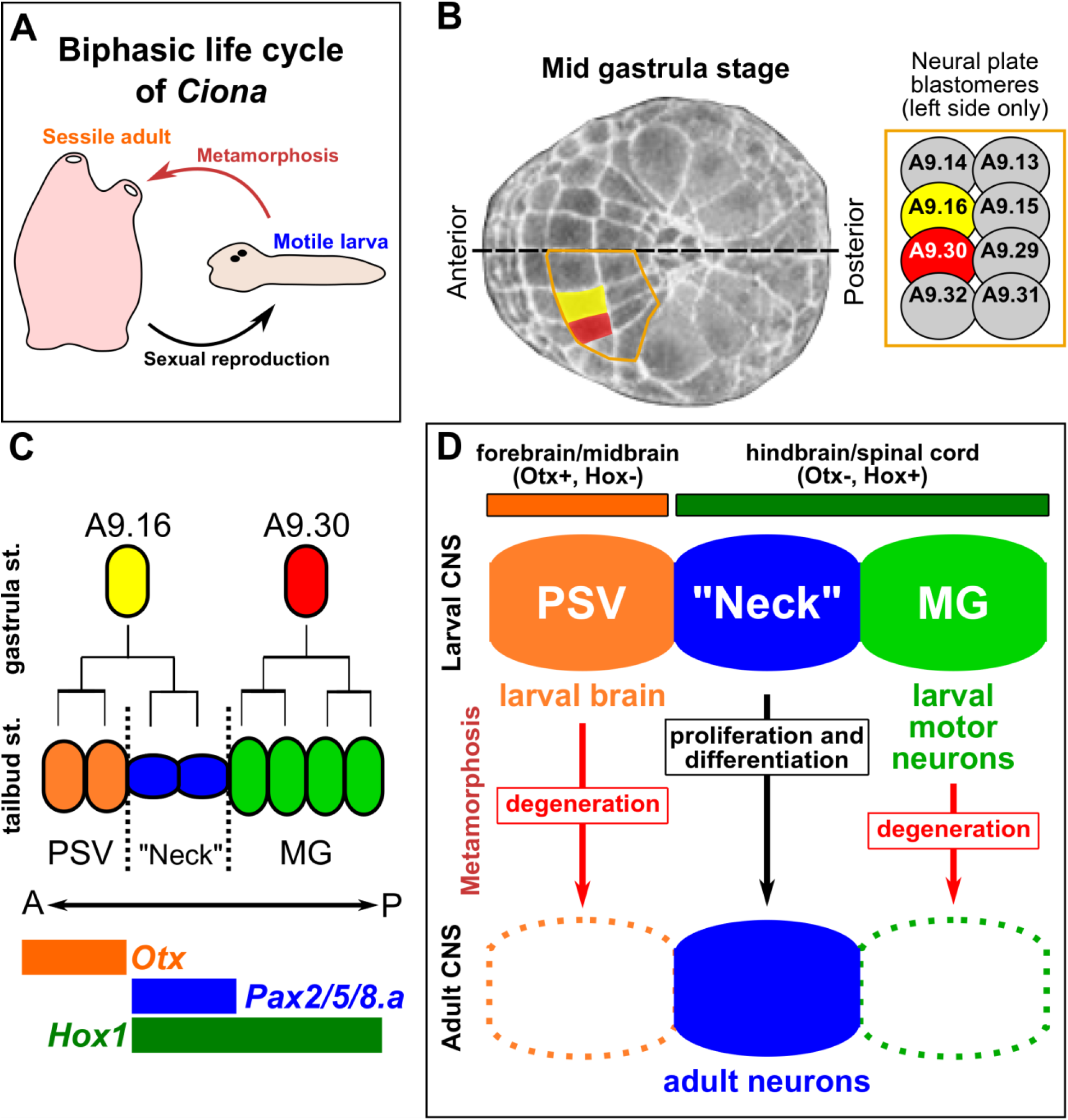
Specification of the “Neck” during *Ciona* embryogenesis. A) A diagram of the biphasic life cycle of *Ciona* and many other tunicates, alternating between a motile, non-feeding larva (pre-metamorphic) and a sessile, filter-feeding juvenile/adult (post-metamorphic). B) Image of mid gastrula-stage *Ciona robusta* embryo (St. 12), adapted from the Tunicanato Database (Hotta et al. 2020), with the A9.16 (yellow) and A9.30 (red) blastomeres false-colored in the neural plate. The left half of the vegetal pole-derived neural plate is outlined in orange, and blastomere identities displayed in the inset at right. Dorsal midline indicated by dashed line. C) Simple diagram of the cell lineages derived from the A9.16 and A9.30 blastomeres giving rise to neurons, photoreceptors, and undifferentiated precursors of the Posterior Sensory Vesicle region (PSV) in orange, the “Neck” in blue, and cells of the Motor Ganglion in green. Anterior(A)-Posterior(P) axis shown as left to right. Colored bars indicate expression domains of conserved forebrain/midbrain regulatory gene *Otx* in orange, rhombospinal (hindbrain/spinal cord) regulatory gene *Hox1* in green, and the Neck/hindbrain marker *Pax2/5/8.a* in blue. D) Diagram summarizing the “traditional” view of the *Ciona* larval central nervous system (CNS), indicating proposed homology of PSV, Neck, and MG compartments to vertebrate central nervous system partitions, based on *Otx* and *Hox* gene expression patterns. According to this view, larval neurons of the PSV/brain and MG are eliminated during metamorphosis, while the Neck contributes neurons to the post-metamorphic, juvenile/adult nervous system.

In this work, we focused on a specific compartment of *Ciona* adult neural precursor cells in the larva called the “Neck” (**Figure 1B,C**). Based on their expression of conserved regulatory genes such as *Pax2/5/8.a, Phox2,* and *Hox1* (**Figure 1C,D**), it has been proposed that the Neck is homologous to part of the vertebrate hindbrain, specifically those cells giving rise to branchiomeric efferents, such as branchiomeric motor neurons(Dufour et al. 2006; Hudson and Yasuo 2021). Although the Neck is specified during embryonic development and shares a close developmental lineage with differentiated larval brain neurons (**Figure 1B-D**), most of it remains in an undifferentiated state until after metamorphosis (Dufour et al. 2006; Imai et al. 2009). Despite their shared origins, larval neurons degenerate during metamorphosis, while neural progenitors in the Neck and in other compartments appear to survive into the adult phase (**Figure 1D**)(Horie et al. 2011; Hozumi et al. 2015). However, the signaling pathways and regulatory networks that direct Neck specification, survival, and differentiation remain uncharacterized.

Here we investigate the molecular mechanisms underlying the specification and patterning of the Neck from embryogenesis to post-metamorphic development. We show that the Neck cells continue to divide throughout embryogenesis and the larval phase, with some of their derivatives differentiating into larval-and adult-specific neurons. We show that Pax2/5/8.a and Phox2 are required for the specification of Neck-derived adult neurons, and that FGF signaling regulates the balance of differentiation and proliferation/survival in the Neck during the transition between larval and adult phases. Thus, we reveal key molecular mechanisms underlying the ability of *Ciona* to generate an entirely separate adult nervous system even as its larval nervous system is largely eliminated.

## Results

### A time-series description of the “Neck” lineage through development and metamorphosis

Presently, the *C. robusta* Neck has been described as 6 ependymal cells per side, thought to be quiescent in nature, and a bilateral pair of differentiated neurons, the so-called “Neck Neurons” (Ryan et al. 2016; Ryan et al. 2017). To establish how the Neck lineage develops between early tailbud and hatched larval stage, we examined Neck cell number, cell type, and morphology using *Pax2/5/8.a* (Oonuma et al. 2021) and *Phox2* (Dufour et al. 2006) fluorescent reporters at hourly intervals, in embryos raised at 20°C (**Figure 2A-J**).

**Figure 2.**
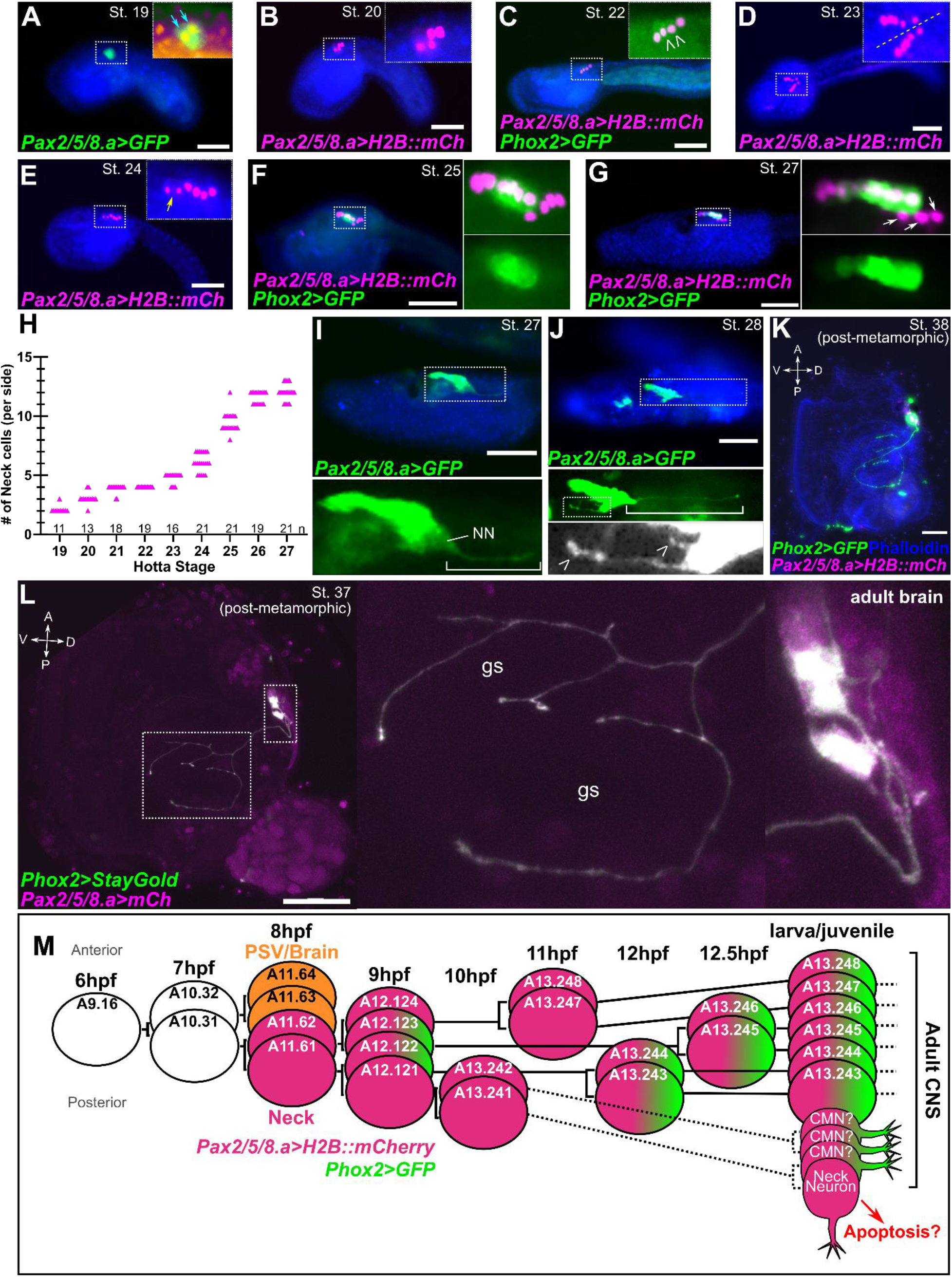
Time-series investigation of cell division and differentiation in the Neck. A-G) A time-series of images of *C. robusta* embryos electroporated with *Pax2/5/8.a* and *Phox2 (Ciona intestinalis/Type B)* reporter plasmids and fixed at different times. Approximate fixation times at 20°C for each stage: St. 19: 8.75 hpf, St. 20: 9 hpf, St. 22: 10 hpf, St. 23: 11 hpf, St. 24: 12 hpf, St. 25: 14 hpf, St. 27: 18 hpf, St. 28: 20 hpf. Insets represent zoomed-in views of areas in dashed rectangles. Nuclei counterstained with Hoechst (blue). Reporter genes used are H2B::mCherry (H2B::mCh) in magenta, and Unc-76::GFP (GFP) in green. Blue arrows in (A) indicating two *Pax2/5/8.a>Unc-76::GFP+* cells. Open arrowheads in (C) and (G) indicate nascent *Phox2(C.intestinalis)>Unc-76::GFP* expression. Yellow arrow in (E) shows cell undergoing mitosis. White arrows in (G) show differentiating neuron nuclei separating from the rest of the Neck, which remains a part of the neural tube epithelium. H) Plot showing the number of *Pax2/5/8.a>H2B::mCherry*+ nuclei counted in embryos from each stage; n = number of embryos examined per stage. I) Stage 27 larva (∼18 hpf) electroporated with *Pax2/5/8.a>Unc-76::GFP,* revealing differentiating Neck Neuron (“NN”) extending its axon posteriorly towards the tail (white bracket in inset). J) Stage 28 larva (∼20 hpf) electroporated with *Pax2/5/8.a>Unc-76::GFP* showing Neck Neuron axon (white bracket, middle inset) and additional axons extending ventrally (open arrowheads in bottom inset), putatively representing early-differentiating post-metamorphic ciliomotor neurons (CMNs). K) Stage 38 (∼96 hpf) post-metamorphic juvenile showing *Pax2/5/8.a>H2B::mCherry+/Phox2(C.robusta)>Unc-76::GFP+* neuronal cell bodies in the adult brain (cerebral ganglion), and GFP+ axons projecting towards the gill slits. Animal counterstained with phalloidin-AlexaFluor 405. See Supplemental Figure 1 for magnified view of cerebral ganglion region and double-labeling with *Pax2/5/8.a>H2B::mCherry.* L) Confocal Z-stack projection of a stage 37 (∼72 hpf) juvenile showing *Pax2/5/8.a>Unc-76::mCherry* (magenta)*, Phox2(C.robusta)>Unc-76::StayGold* (green) CMNs innervating the first two gill slits (gs). Insets showing higher magnification view of gill slits and brain. Merged white (green + magenta) signal due to all GFP signal colocalizing with that of mCherry. A: anterior, P: posterior, D: dorsal, V: ventral. M) Proposed cell lineage and cell division timing for the early Neck. Hypothesized cell divisions displayed with dashed lines. Depicted expression of *Phox2* based on *Phox2>GFP* reporter and not *Phox2* transcript. All scale bars = 50 µm.

At the early tailbud stage (Hotta Stage 19, ∼9 hpf), two *Pax2/5/8.a>H2B::mCherry+* positive cells on either left/right side of the embryo can be observed, corresponding to A11.61 and A11.62 (**Figure 2A**). These two cells continue to divide, forming a single line of four cells on either side by the mid-tailbud stage (St. 20-22, ∼9-10 hpf, **Figure 2B**). By this stage, the middle cells (A12.122 and A12.123) start to express *Phox2>GFP*, whereas the most anterior and posterior cells (A12.121 and A12.124) do not (**Figure 2C**). Starting at Stage 23 (∼11 hpf), the posteriormost cell (A12.121) divides first, (**Figure 2D**), followed by the anteriormost cell (A12.124) dividing around Stage 24 (∼12 hpf, **Figure 2E**). This brings the total number of *Pax2/5/8.a+* cells on either side to six. These six Neck cells have not yet differentiated at this stage and maintain a typical ependymal cell morphology within the neural tube epithelium. Although the cells appear to divide in a stereotyped order, the exact timing of their divisions is slightly variable from embryo to embryo and even between left/right sides in the same individual (**Supplemental Figure 3A**).

During the late tailbud stage (St. 25, ∼14 hpf), the Phox2*+* cells (A12.122 and A12.123) divide, resulting in four cells labeled by *Phox2>GFP* (**Figure 2F**). The posterior*, Phox2*-negative cells also divide around this time and appear to delaminate from the neural tube epithelium, separating from the *Phox2+* cells (**Figure 2F,G**). Beyond stage 26 (larval stages, ∼16 hpf onwards), additional cell divisions are seen among the *Phox2*>GFP*+* cells, as well as the anterior *Phox2*-negative cells. This increases the number of *Pax2/5/8.a+* cell to an average of 12 on each side but the order of divisions and lineage are not easily traced (**Figure 2G**,**H**). Furthermore, *Phox2>GFP* expression expands anteriorly in the lineage during the larval phase but is not detectable in the posterior cells that have delaminated from the neural tube epithelium (**Figure 2G**). The average number of *Pax2/5/8.a+* Neck cells was quantified at each developmental stage and is reported in **Figure 2H**. Based on these data, Neck cells continue to divide into the larval stage, suggesting they are not as quiescent as previously assumed (Imai et al. 2009).

During the larval stages we observed the differentiation of the posteriormost cell of the lineage on either side, likely the “Neck Neuron” as previously described (Ryan et al. 2016; Ryan et al. 2017)(**Figure 2I**). However, we also observed *Pax2/5/8.a>GFP*+ neurites extending from cells just anterior to the Neck Neuron (**Figure 2J**). In early post-metamorphic juveniles (St. 37, 38, ∼72-95 hpf) we observed neurons labeled by *Phox2* and *Pax2/5/8.a* reporters innervating the gill slits (**Figure 2K,L; Supplemental Figure 3B; Supplemental Video 1**), similar to those reported previously (Dufour et al. 2006), and presumably correspond to the cholinergic ciliomotor neurons (CMNs) that control branchial ciliary flow (Jokura et al. 2020). Based on their position, axon trajectory, and expression of *Pax2/5/8.a,* these CMNs are likely to correspond to the early differentiating neurons whose nascent axons can be seen extending towards the future gill slits (**Figure 2J**). Together, these observations suggest that the adult nervous system begins to differentiate before metamorphosis and the degeneration of the larval nervous system. This was surprising given the traditional assumption that larval neurons are eliminated during metamorphosis, while adult neurons arise from set-aside, undifferentiated progenitors. Our results suggest a less clear contrast between pre-and post-metamorphic neurogenesis in tunicates. An updated model of the Neck cell lineage and its derivatives is proposed in **Figure 2M**.

### Pax2/5/8 and Phox2 orthologs are required for the formation of Neck-derived neurons

Having established that the Neck lineage likely gives rise to gill slit-innervating CMNs, we next tested the roles of *Pax2/5/8.a* and *Phox2* in establishing these post-metamorphic neurons. More specifically, we used CRISPR/Cas9 gene editing and gene-specific sgRNAs to disrupt either gene in neural progenitors prior to hatching and metamorphosis. Embryos were electroporated with “negative control” or gene-specific sgRNA expression plasmids, *Sox1/2/3>Cas9::Geminin^N-ter^*plasmid to drive expression of Cas9 in neural progenitors, and a *Phox2>GFP* reporter to label post-metamorphic CMNs. Animals were grown to Stage 37 (∼72 hpf) and imaged to assay the presence of CMNs. In negative control animals, *Phox2>GFP+* CMNs were observed in a majority of animals (**Figure 3A,D**). In contrast, disrupting either *Pax2/5/8.a* or *Phox2* by CRISPR resulted in drastic loss of *Phox2>GFP+* CMNs, as evidenced by lack of axons innervating the gill slits (**Figure 3B-D**). This suggests that Pax2/5/8.a and Phox2 are necessary for the specification and differentiation of post-metamorphic CMNs, supporting their origins from the Neck lineage in which both transcription factors are initially detected.

**Figure 3.**
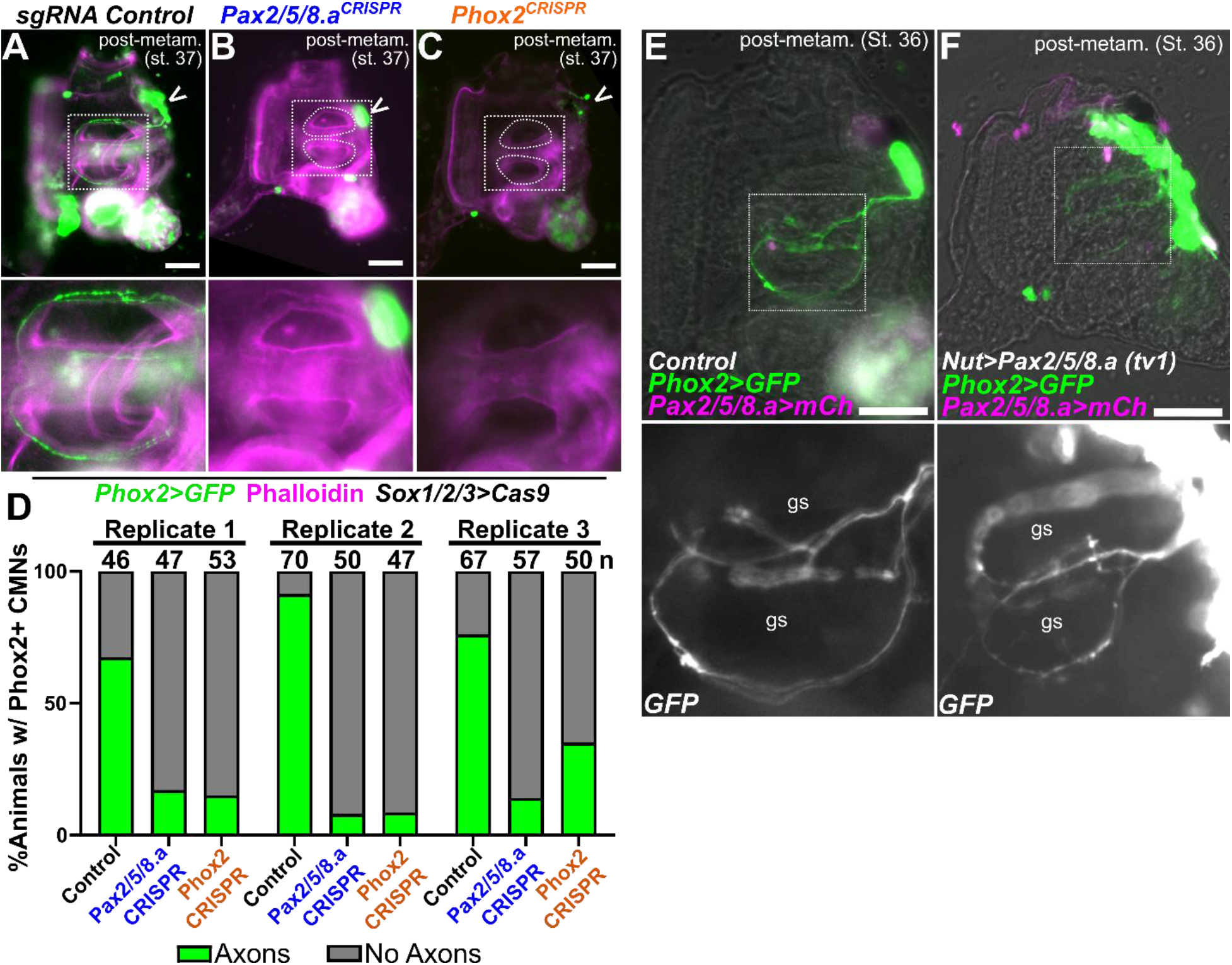
Regulation of adult ciliomotor neuron specification by Pax2/5/8.a and Phox2. A) “Negative control” stage 37 juvenile electroporated with *Sox1/2/3>Cas9::Geminin^N-ter^*and the U6>Control sgRNA expression vector, showing normal pattern of *Phox2 (C.robusta)>Unc-76::GFP* reporter expression in neuronal cell bodies in the brain (open arrowhead) and axons innervating the gill slits (inset). B) Stage 37 juvenile in which neural-specific knockout of *Pax2/5/8.a* by CRISPR/Cas9 has resulted in loss of gill slit-innervating axons (inset), suggesting loss of ciliomotor neurons (CMNs). *Phox2* reporter expression in the brain (open arrowhead) suggests some Phox2+ cells may still be specified. Counterstain in A-C using phalloidin-Alexa Fluor 647. C) Stage 37 juvenile in which neural-specific knockout of *Phox2* has been achieved by CRISPR/Cas9, also resulting in loss of CMN axons innervating the gill slits (inset), and sparse Phox2+ cells remaining in the brain (open arrowhead). D) Scoring of *Phox2+* CMN axon across all three conditions represented in panels A-C, in three independent replicates (n = number of individuals scored in each sample). Individuals were scored for presence or absence of *Phox2>Unc-76::GFP* CMN axons innervating the gill slits. E) Stage 36 juvenile (∼72 hpf) showing electroporated with *Phox2(C.robusta)>Unc-76::GFP* (green) and *Pax2/5/8.a>H2B::mCherry* (magenta nuclei), and a negative control *Nut>lacZ* plasmid. Inset showing gill slits (gs) innervated by CMNs. H) Stage 36 juvenile electroporated with same reporters as in (F), plus *Nut>Pax2/5/8.a (tv1)* to drive expression of Pax2/5/8.a in the entire larval neural tube. *Phox2* reporter is expanded in the resulting juveniles, though there is no noticeable increase in CMN axons innervating the gill slits (gs, inset). All scale bars = 50 µm.

To understand how overexpression of Pax2/5/8.a impacts the developing nervous system, we expressed *Pax2/5/8.a* transcriptional variant 1 (*tv1*) under control of the pan-neuronal Nut driver (*Nut>Pax2/5/8.a*) and examined *Phox2>GFP*+ cells in the Stage 36 juvenile (∼72 hpf). In *Nut>LacZ* negative controls, *Phox2>GFP+* CMNs innervate the early gill slits as expected (**Figure 3E**). In animals overexpressing Pax2/5/8.a, we observed an expansion of Phox2+ cells throughout the juvenile brain region, but no change in the presence or number of gill slit-innervating CMN axons (**Figure 3F**). Together, this suggests that Pax2/5/8.a is sufficient to activate Phox2 expression in the post-metamorphic CNS, but might not be sufficient to impart a CMN identity.

### Characterizing the transcriptional program downstream of Pax2/5/8 by RNAseq

Because Pax2/5/8.a appeared to be a key regulator of Neck identity, we sought to identify its potential downstream transcriptional targets. To do this, we measured global transcriptome changes by bulk RNA sequencing (RNAseq). We used the *Nut* promoter(Shimai et al. 2010) to overexpress both isoforms of *Pax2/5/8.a* (transcript variants “tv1” and “tv2”) throughout the neural tube, and compared whole-embryo transcriptomes of Pax2/5/8.a-overexpression and negative control (no overexpression) embryos at 10 hpf. Differential gene expression analysis revealed the enrichment, or depletion, of transcripts upon overexpression of either Pax2/5/8.a variant (**Figure 4A, Supplemental Table 1**). Previously known Neck markers and Pax2/5/8.a targets (e.g. *Phox2, Gli, Eph.c, FGF9/16/*20) were among the top 600 upregulated genes (out of ∼16,000), with some being considerably higher in ranking (e.g. *Phox2*). In contrast, several known markers of the brain and Motor Ganglion were among the top 50 genes most *downregulated* by Pax2/5/8.a (e.g. *Pax6, Nk6, Neurogenin, BCO).* Correlation between differential gene expression values elicited by the two isoforms was modestly high (Pearson Correlation r = 0.70) (**Supplemental Figure 4**), suggesting high reproducibility and specificity of the Pax2/5/8.a-downstream effects.

**Figure 4.**
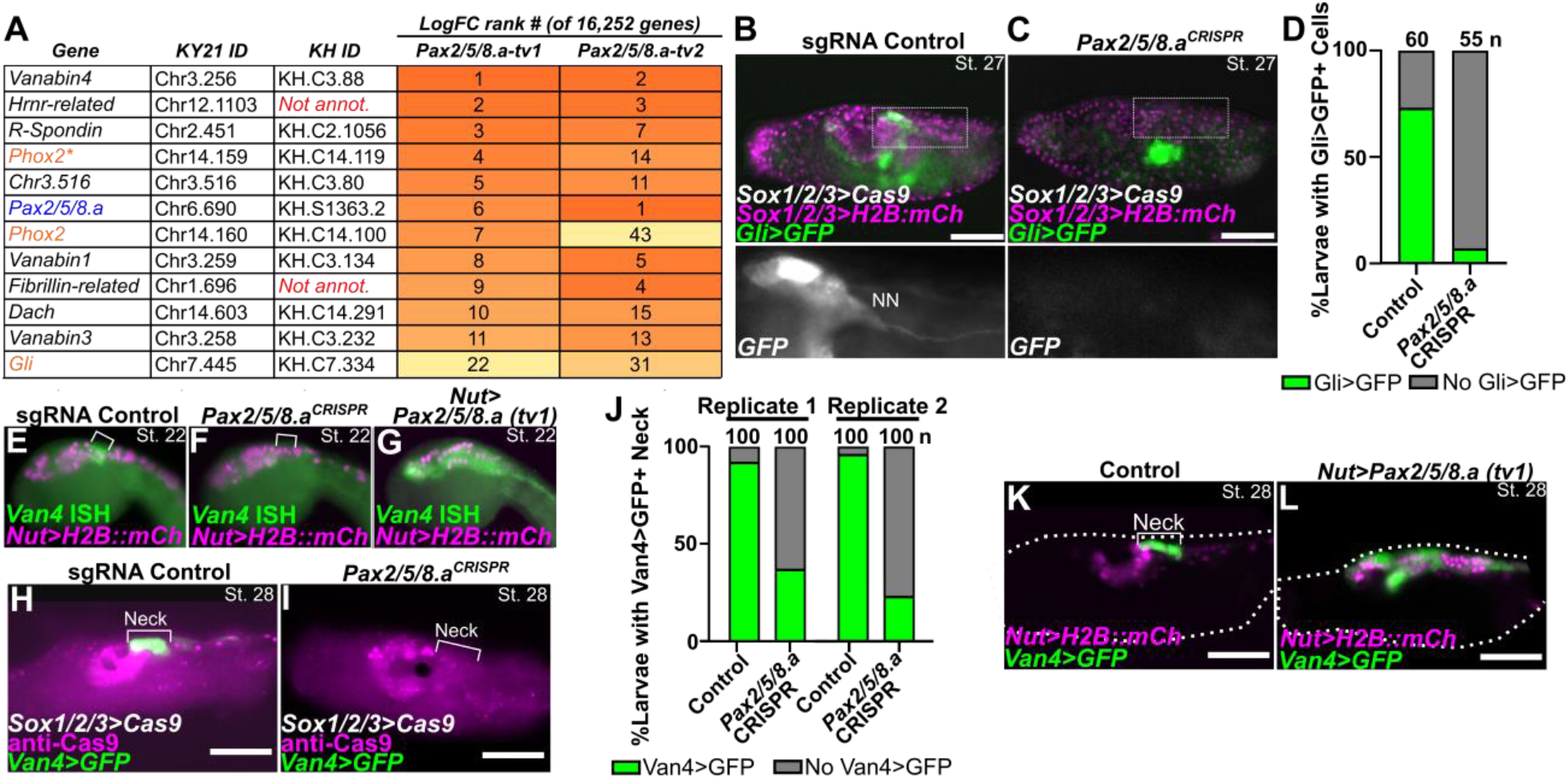
Identifying a Neck-specific gene expression downstream of Pax2/5/8.a. A) Table of rank ordered genes upregulated by Pax2/*5*/8.a overexpression in *Ciona* embryos compared to negative control, as measured by bulk RNAseq. LogFC rank # = ranking of gene when sorting all genes by average Log2 fold-change between the negative control condition (*Nut>lacZ)* and overexpression conditions (*Nut>Pax2/5/8.a, tv1 or tv2).* See text for details. Kyoto 2021 (KY21) and KyotoHoya (KH) gene identification numbers are given for each gene. Genes in orange font are previously known Pax2/5/8.a targets, identified by morpholino knockdown (Imai et al. 2009). Asterisk denotes C-terminal fragment gene model of *Phox2* (see text for details). Differential expression of 16,252 gene models was analyzed in total. B) Negative control animal showing *Gli>Unc76:GFP (Gli>GFP; green)* reporter gene expression in the Neck and Neck neuron (NN). C) CRISPR/Cas9-mediated mutagenesis of *Pax2/5/8.a* results in loss of *Gli>GFP* reporter gene expression in the Neck. *Sox1/2/3* promoter was used to drive expression of Cas9::Geminin^N-ter^ and H2B::mCherry (magenta nuclei) in the ectoderm, including the nervous system. D) Scoring of *Gli+* Neck cells for samples represented in panels B and C. Individuals were scored for presence or absence of *Gli>GFP* cells in the Neck region, posterior to the ocellus and otolith. E) *In situ* mRNA hybridization for Pax2/5/8.a downstream gene *Vanabin4 (Van4)* in negative control embryo (left), showing specific expression in the Neck (bracket) at Stage 22 (10hpf). F) *Pax2/5/8.a* knockout by CRISPR/Cas9 in the nervous system (using *Sox1/2/3>Cas9)* results in loss of *Van4* mRNA *in situ* signal in the Neck. G) Overexpression of *Pax2/5/8.a (transcript variant 1)* in throughout the neural tube (*Nut>Pax2/5/8.a tv1)* results in widespread, ectopic expression of *Van4*, confirming this gene as being downstream of Pax2/5/8.a as indicated by RNAseq. All embryos electroporated with *Nut>H2B::mCherry* (magenta nuclei). (H) *Van4>Unc-76::GFP* reporter expression (green) also specifically labels the Neck (bracket) in Stage 28 negative control larvae (∼20 hpf). I) Knockout of *Pax2/5/8.a* in the nervous system results in loss of *Van4* reporter expression. Cas9 protein visualized by immunostaining (magenta). J) Scoring of *Van4>Unc-76::GFP* reporter expression in *Pax2/5/8.a* CRISPR larvae represented in H and I, showing dramatic reduction in frequency of *Van4+* larvae in *Pax2/5/8.a* CRISPR larvae compared to negative control larvae, in two independent replicates. K) Stage 28 larva electroporated with *Nut>H2B::mCherry* (magenta nuclei), *Van4>Unc-76::GFP* (green) reporters, and *Nut>LacZ* control. L) Larva electroporated with same reporters as in (K), together with *Nut>Pax2/5/8.a tv1,* showing expansion of *Van4* reporter expression throughout the neural tube. In D and J, n = number of individuals scored in each sample. All scale bars = 50 µm.

One gene previously known to be downstream of Pax2/5/8.a in the Neck, *Gli*(Imai et al. 2009), was among the top 31 upregulated genes using either Pax2/5/8.a isoform (**Figure 4A**). Indeed, CRISPR/Cas9-mediated knockout of *Pax2/5/8.a* largely abolished *Gli>GFP* reporter expression in the Neck, confirming that it is downstream of Pax2/5/8.a (**Figure 4B-D**). Following the same approach, we were able to validate a novel putative target of Pax2/5/8.a, the gene *Vanabin4* (*Van4,* gene IDs *KH.C3.88/KY21.Chr3.256*). *Van4* encodes a tunicate-specific vanadium-binding protein and was among the top 1-2 upregulated genes by both isoforms of Pax2/5/8.a (**Figure 4A**). *In situ* mRNA hybridization revealed expression of *Van4* specifically in the Neck (**Figure 4E**). *Van4* expression was lost upon *Pax2/5/8.a* CRISPR knockout (**Figure 4F**), and expanded throughout the neural tube upon overexpression of Pax2/5/8.a (**Figure 4G**). *Van4>GFP* reporter expression in the Neck was similarly lost upon *Pax2/5/8.a* CRISPR knockout (**Figure 4H-J**). Taken together, these data reveal a transcriptional program for Neck specification downstream of Pax2/5/8.a in *Ciona*.

### Correcting the C. robusta Phox2 gene model

During analysis of RNA sequencing data, we observed upregulation of *Phox2* (*KY21.Chr14.160*) and neighboring gene *KY21.Chr14.159* by Pax2/5/8.a overexpression. On closer inspection of our bulk RNA sequencing, we found that reads spanned exons covering both gene models, and further revealed two cryptic exons (exon 1 and exon 5)(**Supplemental Figure 5A,B**). A corrected, consolidated gene model (**Supplemental Figure 5C**) was supported by sequencing a full-length Phox2 cDNA clone spanning all 7 exons, cloned from 17 hpf larvae (**Supplemental Figure 5D**), and by protein sequence alignment with predicted Phox2 protein models from other tunicate species (**Supplemental Figure 5E**). Finally, mRNA *in situ* hybridization using probes designed for both *Chr14.160* and *Chr14.159* gene models revealed identical expression patterns specific to the Neck, further supporting the finding that these are indeed the same gene (**Supplemental Figure 5F,G**). The function of the extended C-terminus (which is divergent relative to that of vertebrate Phox2 family members) is unknown, but will likely be important for future studies on transcriptional regulation by Phox2.

### Characterization of Neck cell transcriptomes in single-cell RNAseq data

To further characterize the molecular profiles of Neck cells, we re-analyzed published whole-embryo single-cell RNA sequencing (scRNAseq) data obtained at the mid-tailbud II stage(Cao et al. 2019). This revealed a single cluster enriched for *Pax2/5/8.a* reads (cluster 25, **Figure 5A, Supplemental Figure 2A,B, Supplemental Table 2**). Re-clustering performed only on cluster 25 cells revealed two distinct subclusters, with *Pax2/5/8.a* enriched in one subcluster (subcluster 0) and depleted in the other (subcluster 1)(**Figure 5B, Supplemental Figure 2C**). Double *in situ* hybridization using probes for *Pax2/5/8.a* and a top subcluster 1 marker *Crls1 (KH.C11.724)* confirmed that subcluster 0 represents the Neck, while subcluster 1 appears to represent larval brain neurons and photoreceptors just anterior to the Neck (**Figure 5C**). Other markers enriched in subcluster 0 cells further confirmed their identity as Neck cells, showing substantial overlap with the top genes upregulated by Pax2/5/8.a overexpression as measured by our bulk RNAseq experiment above (**Figure 5D,E** **Supplemental Table 3**). Correlation between enrichment/depletion in the Neck by scRNAseq and average upregulation/downregulation by Pax2/5/8.a was modest (Pearson correlation = 0.43; Spearman’s rank correlation = 0.46)(**Supplemental Table 4**). We observed notable exceptions, such as *Hox1,* which is highly expressed in the Neck but was not upregulated by Pax2/5/8.a (**Figure 5D,E**). This suggests that Pax2/5/8.a is not sufficient to activate the entirety of the transcriptional program of the Neck, and that some important Neck regulators might be expressed in parallel to, not downstream of, Pax2/5/8.a. In contrast, the top markers enriched in the “brain” subcluster (subcluster 1) were among those transcripts most highly depleted by Pax2/5/8.a overexpression, suggesting that Pax2/5/8.a is also instrumental for repressing larval brain neuron/photoreceptor identity.

**Figure 5.**
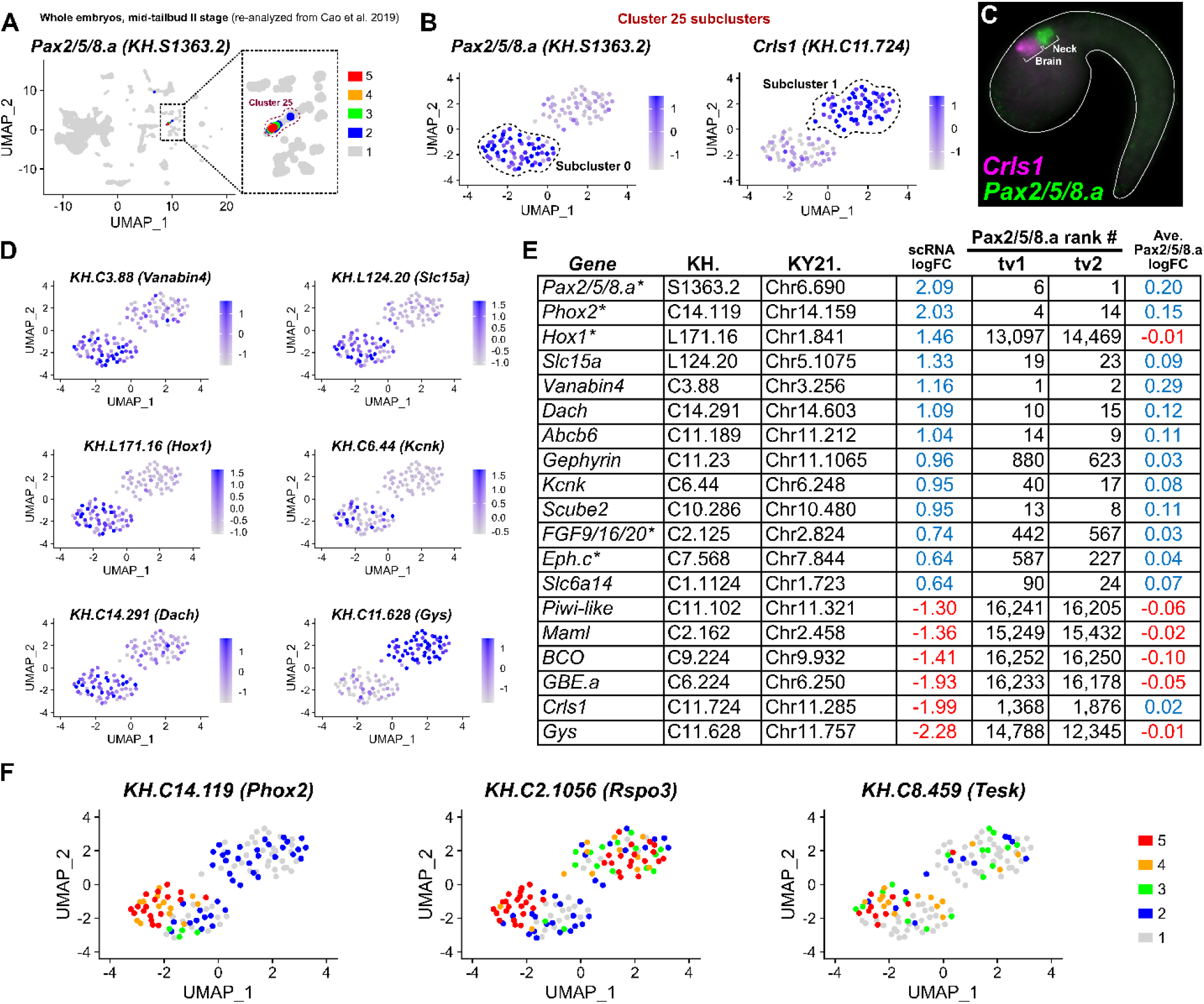
Analyzing Neck-specific gene expression in single-cell RNAseq data. A) Re-analysis of published whole-embryo *C. robusta* single-cell RNA sequencing (scRNAseq) data from Cao et al. (2019) (Cao et al. 2019) revealed a cluster of cells (Cluster 25) enriched for Neck marker *Pax2/5/8.a* (magnified view of boxed area in inset). Differential gene expression “FeaturePlot” color-coded as measured by “RNA” assay in Seurat. B) Differential gene expression as measured by “integrated” assay of reclustered cells from Cluster 25 showing enrichment of *Pax2/5/8.a* reads in Subcluster “0” relative to subcluster “1”, and enrichment of *Crls1* in subcluster “1” relative to subcluster “0”. C) Two-color double mRNA *in situ* hybridization for *Crls1* (magenta) and *Pax2/5/8.a* (green) in a Stage-22 embryo confirm that subcluster “0” represents the Neck and subcluster “1” represents brain/posterior sensory vesicle cells just anterior to the Neck. D) Differential gene expression FeaturePlots showing enrichment of various genes in the Neck (subcluster 0) relative to the brain (subcluster 1), or vice-versa (e.g. *Glycogen synthase, or Gys).* All plots generated by “integrated” assay in Seurat. E) Table of genes comparing enrichment (blue font) or depletion (red font) in subcluster “0” cells to upregulation upon overexpression of *Pax2/5/8.a* transcript variants 1 and 2. “Ave. Pax2/5/8/a logFC” = average tv1 and tv2 logFC values as compared to a negative control. Asterisks denote previously known Neck marker genes. *Hox1* is a rare exception of a gene that is enriched in the Neck by scRNAseq but is not upregulated by Pax2/5/8.a. F) Differential gene expression FeaturePlots (5-color scale, “integrated” assay) showing higher expression of *Phox2, Rspo3,* and *Tesk* in roughly half of subcluster 0, recapitulating the distinction between *Phox2+* middle cells and flanking Phox2-negative cells seen by reporter assays (Figure 2).

As expected, *Phox2* expression (represented by the KH.C14.119/KY21.Chr14.159 gene model, due to the 3’ bias of 10X Genomics system) was unevenly distributed within subcluster 0. *Phox2* was expressed more highly in one half of the subcluster than in the other half (**Figure 5F**), recapitulating its expression in a subset of Neck cells as seen by *Phox2>GFP* expression. Additional genes showed a similar distribution in Neck subcluster (e.g. *Rspo3, Tesk,* **Figure 5F, Supplemental Figure 2D**), further confirming the distinction between proliferating/undifferentiated cells in the anterior Neck and differentiating Neck-derived neurons in the posterior.

### Ephrin/Eph and FGF signaling regulate the balance between proliferation and differentiation

FGF/MAPK signaling plays numerous, recurring roles in the development of the *Ciona* larval nervous system (Davidson et al. 2006; Stolfi et al. 2011; Haupaix et al. 2013; Razy-Krajka et al. 2018). In fact, FGF8/17/18 from the neighboring A9.30 cell lineage is required for expression of *Pax2/5/8.a,* which in turn activates expression of *FGF9/16/20* in the Neck (**Figure 6A**)(Imai et al. 2009). Furthermore, Ephrin/Eph signaling antagonizes FGF/MAPK signaling to provide crucial spatial information for FGF/MAPK-dependent patterning and cell fate choice in the nervous system, including in the Motor Ganglion (MG), in which downregulation of FGF/MEK promotes cell cycle exit and neuronal differentiation (Stolfi et al. 2011; Haupaix et al. 2013). Expression of *FGF9/16/20* and *Eph.c* (formerly *Eph3*) in the Neck (**Figure 6A,B**), and *EphrinA.b* and *EphrinA.d* (**Figure 6C,D**) in the cells of the brain and MG abutting the Neck suggested that FGF and Ephrin/Eph signaling might be key to regulating cell fate and neuronal differentiation in the Neck as well.

**Figure 6.**
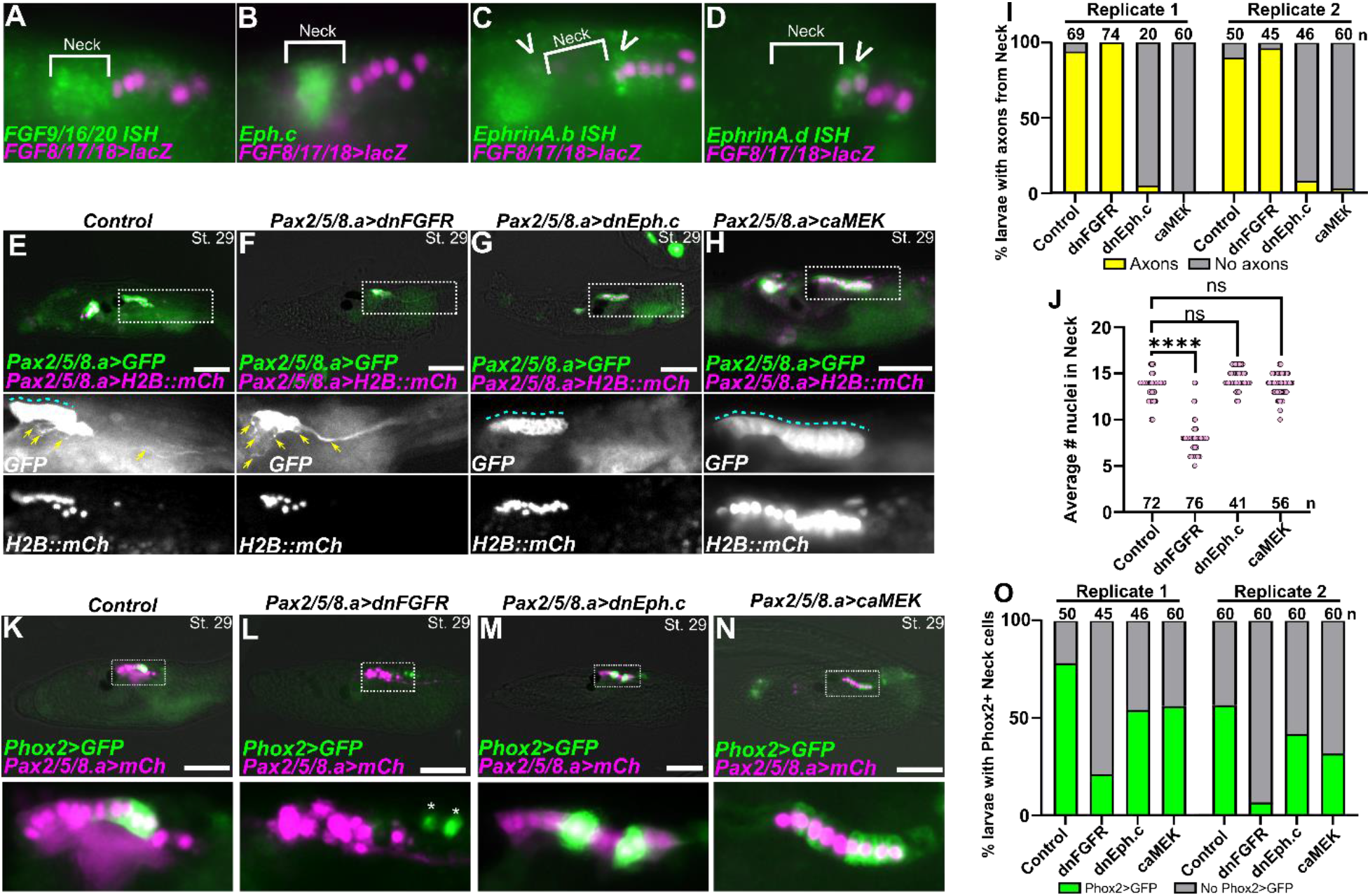
Ephrin/Eph and FGF/MEK signaling pathways regulate differentiation in the Neck. A) *In situ* hybridization (ISH) for *FGF9/16/20* (green), showing expression in the Neck at stage 22. B) ISH for *Eph.c* (green), showing expression in the Neck (St. 22), more strongly in middle cells. C) ISH for *EphrinA.b* (green), showing expression in the anterior cells of the A9.30 lineage as well as the brain region (open arrowhead), both abutting the limits of the Neck. D) ISH for *EphrinA.d* (green), showing expression in the anterior cells of the A9.30 lineage (open arrowhead). In panels A-D, neighboring A9.30 lineage is marked by *FGF8/17/18>LacZ,* revealed by β–galactosidase immunostaining (magenta nuclei). E) Negative control Stage 29 larva (∼21 hpf) electroporated with *Pax2/5/8.a>LacZ* control, *Pax2/5/8.a>Unc-76::GFP* (green), and *Pax2/5/8.a>H2B::mCherry* (magenta nuclei) showing the Neck giving rise to both undifferentiated cells forming an epithelium in the neural tube (blue dashed line) as well as differentiating neurons and extending axons (yellow arrows). F) Larva electroporated with *Pax2/5/8.a>dnFGFR,* showing loss of epithelial structure in the Neck and supernumerary *Pax2/5/8.a+* axons. G) Larva electroporated with *Pax2/5/8.a>dnEph.c,* showing expansion of the the undifferentiated neuroepithelium and loss of axons emanating from the Neck. H) Larva electroporated with *Pax2/5/8.a>caMEK,* which also expanded the neuroepithelial state and suppressed neuronal differentiation and axon growth. I) Scoring of larvae represented in panels E-H, showing almost complete loss of Neck-derived axons in larvae expressing dnEph.c or caMEK in the Neck, across two replicates. J) Plot showing average number of *Pax2/5/8+* nuclei counted in larvae represented in panels E-H, showing significantly fewer nuclei in the dnFGFR condition. K) Negative control larva electroporated with *Pax2/5/8.a>LacZ* control, with *Pax2/5/8.a>Unc76:mCh* and *Pax2/5/8.a>H2b:mCh* (labeled as *Pax2/5/8.a>mCh),* showing *Phox2(C.robusta)>Unc-76::GFP* expression in a subset of Neck cells (green). L) Overexpression of dnFGFR in the Neck abolished *Phox2>GFP* reporter expression. M) Overexpression of dnEph.c expands *Phox2* expression in a variable manner. N) caMEK overexpression is similar to that of dnEph.c, resulting in variable expansion of *Phox2* reporter expression. O) *Phox2>Unc-76::GFP* expression was scored in larvae represented in panels K-N, across two independent replicates. While dnFGFR results in loss of *Phox2* reporter expression, dnEph.c and caMEK do not. In panels I, J, and O, n = number of individuals scored in each sample. All scale bars = 50 µm.

To test the role of FGF and Ephrin signaling in the Neck, we used the *Pax2/5/8.a* promoter to overexpress dominant-negative FGF receptor (dnFGFR), dominant negative Eph.c receptor (dnEph.c), or a constitutively-active form of the MAPK kinase MEK (MEK^S220E,S216D^, also called caMEK)(Davidson et al. 2006; Picco et al. 2007; Shi and Levine 2008; Razy-Krajka et al. 2018). We examined the Neck at the late swimming larval stage (St. 29), when there are ∼8 anterior putatively undifferentiated, neuroepithelial cells and ∼4 posterior, differentiating neurons in control larvae (**Figure 6E,I,J**). In larvae expressing the dnFGFR, we observed fewer cells overall in the Neck, and undifferentiated cells appeared to be replaced by supernumerary neurons, as evidenced by loss of epithelial organization and excess axon outgrowth (**Figure 6F,I,J**). This contrasted starkly with larvae expressing the dnEph.c, in which we observed a near-complete loss of differentiated neurons, most clearly evidenced by a distinct absence of the posterior Neck Neuron axon extending towards the tail (**Figure 6G,I,J**). Instead, the entire Neck took on the morphology of undifferentiated neural precursors in a tightly-packed neuroepithelium. This was phenocopied by caMEK, which constitutively activates the FGF/MAPK pathway in all cells (**Figure 6H,I,J**). We observed these phenotypes, consistently across two biological replicates (**Figure 6I,J**). Taken together, these results suggest that Ephrin/Eph-mediated suppression of FGF/MAPK signaling is sufficient and necessary for neuronal differentiation in the Neck, and that sustained FGF/MAPK signaling promotes an undifferentiated, neuroepithelial state instead.

We also examined the expression of *Phox2>GFP* in the Neck, which is normally expressed by the ependymal-like cells in the middle part of the Neck (**Figure 6K**). As expected, we observed a loss of Phox2+ Neck cells in larvae electroporated with *Pax2/5/8.a>dnFGFR* (**Figure 6L,O**). However, neither *Pax2/5/8.a>dnEph.c* nor *Pax2/5/8.a>caMEK* abolished *Phox2>GFP* expression but rather made its spatial distribution within the Neck inconsistent (**Figure 6M-O**). This suggests that sustained FGF/MEK signaling is permissive but not sufficient for *Phox2* activation in the Neck.

Because our data had revealed upregulation of genes encoding the BMP antagonist Noggin and the Hedgehog (Hh) pathway effector Gli, downstream of Pax2/5/8.a (**Figure 4, Supplemental Table 4**), we also tested the effects of perturbing these pathways. No significant effect on Neck morphogenesis was observed upon overexpression of dominant-negative or constitutive BMP receptors (**Supplemental Figure 6A-D**) or CRISPR/Cas9-mediated knockout of *Gli* (**Supplemental Figure 6E-G**). While these results do not rule out a role for BMP and/ Hh signaling pathways in the development of the Neck or Neck-derived neurons, they do not impact the balance of differentiation and proliferation like Ephrin and FGF do.

Finally, because disruption of Ephrin/FGF/MAPK signaling significantly impacted cell proliferation and differentiation in larvae, we examined how these perturbations affected the development of CMNs in post-metamorphic juveniles. We observed a loss of *Pax2/5/8.a>GFP*+ CMN axons in juveniles that had expressed the dnFGFR receptor in the Neck at the larval stage (**Figure 7A,B**), but not in animals that expressed dnEph.c or caMEK (**Figure 7C,D**). We conclude that the supernumerary, differentiating neurons seen in the larvae generated by *Pax2/5/8.a>dnFGFR* do not survive metamorphosis, failing to give rise to fully differentiated CMNs in the juvenile. In contrast, undifferentiated neuroepithelial cells elicited by dnEph.c or caMEK overexpression still retain the potential to differentiate into CMNs after metamorphosis. Taken together, these data suggest that the balance between neuronal differentiation in the larva and survival of neuronal precursors set aside for the adult CNS likely depends on a careful balance of FGF signaling prior to metamorphosis.

**Figure 7.**
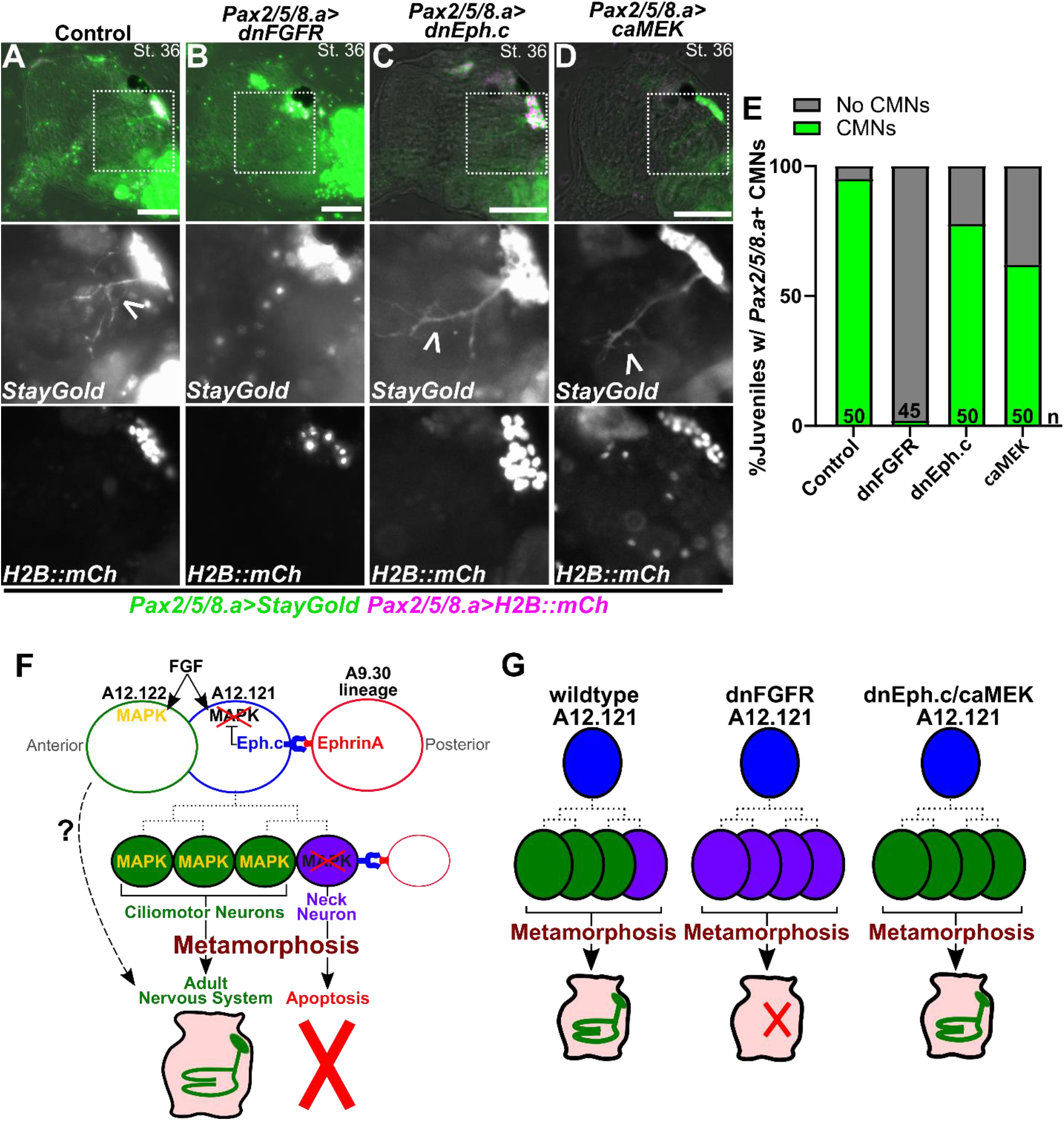
FGF signaling is required for survival of CMN precursors through metamorphosis. A-D) Stage 36 juveniles (∼72 hpf) electroporated with *Pax2/5/8.a>Unc-76::StayGold* (*Pax2/5/8.a>StayGold*; green) and *Pax2/5/8.a>H2B::mCherry* (magenta), showing growing ciliomotor (CMN) axons (open arrowhead) in the process of innervating the gill slits. A) Negative controls expressing *Pax2/5/8.a>LacZ* show normal CMNs. B) *Pax2/5/8.a>dnFGFR* eliminates CMN axons, suggesting maintenance of FGF/MAPK signaling is important for specification of CMNs and survival through metamorphosis. C) Juveniles expressing dnEph.c still give rise to CMNs (open arrowhead) during metamorphosis, indicating an ability to recover from earlier suppression of differentiation in the larva (see Figure 7). D) Juvenile expressing caMEK also recover, growing CMN axons (open arrowhead) during metamorphosis. E) Scoring of juveniles represented by panels A-D, showing near total loss of CMNs in juveniles derived from larvae expressing dnFGFR in the Neck. F) Diagram explaining model for Ephrin/Eph-mediated suppression of FGF/MAPK signaling in the posteriormost cell of the Neck, which becomes the Neck Neuron, and hypothesized cell lineage from A12.121 (dashed lines). More anterior cells escape Ephrin signaling at different times, giving rise to precursors of the adult nervous system including CMNs, while the Neck Neuron does not appear to survive metamorphosis. The uncertain contribution of A12.122 derivatives to the adult nervous system indicated by dashed arrow. G) Diagram explaining the different manipulations tested and their effects on CMN formation in juveniles. In wildtype animals, the Neck Neuron (lavender) does not survive metamorphosis, and more anterior CMN precursors differentiate during late larval/early juvenile stages. In *Pax2/5/8.a>dnFGFR-*electroporated animals, the entire Neck is converted to Neck Neuron-like neurons, which are all eliminated during metamorphosis. In individuals electoporated with *Pax2/5/8.a>dnEph.c* or *Pax2/5/8.a>caMEK,* sustained FGF signaling temporarily suppresses neuronal differentiation in the Neck during larval stages, but the cells eventually “recover” during metamorphosis to give rise to CMNs.

## Discussion

Herein, we have described the development of the *Ciona* “Neck” beyond the tailbud stage, through metamorphosis and post-metamorphic neuronal differentiation. Despite reports of the Neck cells being quiescent in the larva, we observe ongoing proliferation and precocious ciliomotor neuron (CMN) differentiation throughout the larval stage. We also show for the first time that transcription factors *Pax2/5/8.a* and *Phox2* are essential for the formation of adult (post-metamorphic) CNS neurons. Through RNA sequencing we have identified the Neck transcriptional program downstream of *Pax2/5/8.a,* revealing candidate target genes of Pax2/5/8.a that might be important for post-metamorphic neurodevelopment. Finally, although we did not identify an obvious role for either the BMP or Hedgehog pathways, we establish that Ephrin and FGF signaling are responsible for patterning the Neck into differentiated and undifferentiated compartments, with serious consequences for adult CNS formation if disturbed.

Across many species, *Pax2*/5/8 homologues are midbrain/hindbrain regulators suggesting they are part of an ancestral gene regulatory network (Bridi et al. 2020; Schuster and Hirth 2023). In vertebrates, Phox2 proteins regulate the development of various motor neurons of the head, which are also cholinergic (Pattyn et al. 2000; Mazzoni et al. 2013; Curto et al. 2015). For instance, PHOX2A (along with PAX2/5) establishes oculomotor and trochlear cranial motor neurons at the isthmic organizer boundary between the midbrain and hindbrain (Deng et al. 2011; Fritzsch 2023), while loss of *Phox2b* ablates cranial branchial neurons and visceral motor neurons of the hindbrain in mouse (Pattyn et al. 1999; Pattyn et al. 2000). Also in mouse, Phox2a promotes cranial motor neuron fate over spinal cord motor neuron fate (Mazzoni et al. 2013). In *Ciona*, *Pax2/5/8.a+/Phox2+* Neck cells give rise to post-metamorphic neurons that innervate pharyngeal gill slits and possibly branchiomeric siphon muscles of the adult later on (Wada et al. 1998; Dufour et al. 2006; Hozumi et al. 2015). In contrast, *Ciona Phox2* is not expressed in larval motor neurons, which innervate (paraxial) muscles of the tail, like vertebrate spinal cord motor neurons. Here we show that *Pax2/5/8.a* and *Phox2* are required for the formation of Neck-derived cholinergic CMNs in post-metamorphic *Ciona.* Taken together, these findings suggest that a conserved transcriptional program for cholinergic branchiomeric efferent specification and differentiation might have evolved in the last common ancestor of tunicates and vertebrates, predating the origin of cranial nerves and jaw muscles.

Like many tunicate species, *Ciona* have a biphasic lifecycle; a motile, non-feeding larval stage and a sessile, filter-feeding juvenile/adult stage. It has been traditionally assumed that the larval and adult life stages are separated by a total overhaul during metamorphosis, with the adult CNS completely replacing the larval CNS, thanks to quiescent neural progenitors set aside for the adult nervous system (Horie et al. 2011; Hozumi et al. 2015). However, our observations suggest that the larval-adult boundary might not be as sharp. Instead, we observe precocious differentiation of Neck-derived CMNs that project their axons ventrally and anteriorly during the late larval stage (**Figure 2J**), several hours before tail absorption and body axis rotation (Hotta et al. 2020). Thus, it appears that these particular neurons are already post-mitotic and differentiating during larval phase (Stage 28) and not during the post-metamorphic juvenile stage as previously thought. In juveniles and adults, CMNs have been shown to use cholinergic neurotransmission to regulate ciliary flow in the gill slit epithelium, which generates the water flow that brings suspended food particles into the mouth (Petersen et al. 1999; Jokura et al. 2020). Therefore, the early, pre-metamorphic differentiation of these neurons may be key for the rapid onset of feeding behavior and further growth after metamorphosis.

It is unclear if the Neck also gives rise to additional neurons of the juvenile/adult. Although we and others have seen Phox2+ neurons that are not innervating the gill slits (Dufour et al. 2006), one cannot exclude the possibility that new Phox2+ cells arise from other compartments after metamorphosis. Moreover, as the branchial sack continues to grow and form new gill slits, these structures will also require neural innervation of Phox2+ cells (Dufour et al. 2006). Whether this requires ongoing neurogenesis of Neck-derived Phox2+ cells or additional sources of CMN progenitors is not known. One possibility is that the anteriormost cells of the Neck that activate *Phox2>GFP* later in larval development might contribute to later-differentiating CMNs. It is also unclear why activation of *Phox2* expression appears initially in cells that are not undergoing differentiation yet (**Figure 2G**). It will be important to identify the precise function and transcriptional targets of Phox2 to understand the temporal dynamics of adult neurogenesis in *Ciona*.

In addition to post-metamorphic CMNs, the Neck also gives rise to the Neck Neuron (NN), which here we show is the posteriormost cell of the Neck lineage on either side of the larva, forming a bilaterally symmetric left-right pair of neurons. The NN has been previously described at the early larval stage in the *C. intestinalis* whole-larva connectome studies (Ryan et al. 2016; Ryan and Meinertzhagen 2019). The function of the NN remains unknown, but the connectome described it as receiving synaptic inputs from ascending MG interneuons (AMG neurons) and synapsing primarily onto the basement membrane (Ryan et al. 2016). However, NNs might not be fully differentiated in early larvae, and it is possible they form additional connections later in larval development. Indeed, we noticed that the axon of the NN continues to extend posteriorly during the swimming larval phase and can be observed exiting the trunk and entering the proximal portion of the tail prior to settlement and metamorphosis. The NNs likely do not survive metamorphosis as we rarely observed them in post-metamorphic juveniles. Interestingly, *Pax2/5/8.a>dnFGFR* converted the entire Neck lineage into neurons that did not appear to survive metamorphosis (**Figure 7B**). Because of this, we believe that the supernumerary neurons generated by dnFGFR represent NN-like neurons that do not persist to the adult stage. Although the supernumerary axons in the dnFGFR condition did not project towards the tail like the NNs, this may be due to disrupted neuronal polarity and axon growth mechanisms. Consistent with this hypothesis, we have previously observed supernumerary motor ganglion neurons aberrantly project away from the tail(Stolfi et al. 2011). Future studies will be needed to identify regulatory differences between CMNs and NNs and to investigate what factors determine whether a neuron survives through, or perishes during, metamorphosis despite their shared lineage history. For instance, it is not yet clear if sustained FGF signaling in neural progenitors is inherently pro-survival, or if FGF downregulation in the Neck is simply a molecular switch for the specification of NNs, which may be pre-programmed to degenerate during metamorphosis.

It is possible that certain post-metamorphic cell survival factors are downstream of *Pax2/5/8.a*, including those we identified here using RNAseq approaches. One interesting candidate is *Vanabin4* (*Van4*), which encodes a vanadium-binding protein and was the most highly upregulated target by overexpression of Pax2/5/8.a. The biological function of vanadium or vanabins has remained elusive, and it is not clear what the role of Van4 might be in the Neck (Ueki et al. 2015). As adults, many tunicate species accumulate high levels of intracellular vanadium, but not all species have vanabins and so their purpose remains unknown (Ueki et al. 2003). One theory is that adults accumulate vanadium as a deterrent for predation (Odate and Pawlik 2007), but this alone cannot explain the high expression of vanabins in the Neck. It is possible that vanadium provides a protective effect against oxidative stress (Tripathi et al. 2018), which would promote or enhance survival during metamorphosis. Future studies will examine the role of vanabins in *Ciona* and other marine organisms that also accumulate this transition metal (Thompson et al. 2018; Mendonca et al. 2023)

## Methods

### Ciona handling, fixation, staining, and imaging

Adult *Ciona robusta* (*intestinalis* Type A) were collected from San Diego, CA (M-REP). Dechorionated zygotes were generated and electroporated as previously described(Christiaen et al. 2009a; Christiaen et al. 2009b). Embryos were raised in artificial sea water at 20°C. Animals raised beyond 24 hours post-fertilization (hpf) were moved to new agarose coated plates with fresh artificial sea water +1.0% Penicillin-Streptomycin (Gibco) daily. Post-metamorphic animals were paralyzed with L-menthol prior to fixation, as previously described(Osugi et al. 2020). Staging is based on TUNICANATO database(Hotta et al. 2020). Cell lineage nomenclature is based on Conkin(Conklin 1905; Nicol and Meinertzhagen 1991). All sequences of plasmids, probes, and sgRNAs not previously published can be found in the **Supplement**.

Sample processing for fluorescence and immunostaining were performed as previously described (Beh et al. 2007; Ikuta and Saiga 2007; Stolfi et al. 2011). Unc-76-tagged GFP and mCherry (Dynes and Ngai 1998) were used to improve labelling of cell bodies and axons, instead of GFP/mCherry alone. An Unc-76-tagged StayGold green fluorescent protein was also designed and used for its improved signal longevity(Hirano et al. 2022). Immunolabeling of Cas9 protein by monoclonal anti-mouse Cas9 antibody at 1:500 (4G10; Diagenode) was blocked in PBS Super Block (37580; ThermoFisher) and visualized with goat anti-mouse AlexaFluor 568 (Invitrogen). Juvenile animals stained with phalloidin-AlexaFluor 405 or 647 (ThermoFisher; A30104, A22287). *In situ* hybridization coupled to immunostaining was performed as previously described(Beh et al. 2007), using TSA Plus amplification kits (Akoya Biosciences) and mouse anti-β-galactosidase (Promega #Z378, 1:1000) or rabbit anti-mCherry (BioVision, accession number ACY24904, 1:500) primary antibodies. Two-color (Fluorescein + Cy3) double *in situ* hybridization was performed using TSA Plus amplification kits as previously described(Ikuta and Saiga 2007; Stolfi et al. 2011). Probe template sequences can be found in the Supplemental Sequences file. All standard images were captured on a Leica epifluorescence compound microscope. Confocal images were captured on a Nikon AX R with NSPARC and processed in ImageJ (1.54d). Phenotypes were quantified on a Leica DMi8 or DMIL LED inverted epifluorescence microscope. Biological replicates are presented in side-by-side graphs generated in Prism (9.5.1). Cell count data analyzed by one-way ANOVA and Dunnett’s test of multiple comparisons.

### CRISPR/Cas9 sgRNA design and validation

Single-chain guide RNAs (sgRNAs) were designed using CRISPOR (http://crispor.tefor.net/) and vectors were constructed as previously described (Haeussler et al. 2016; Gandhi et al. 2018) or custom synthesized and cloned (Twist Bioscience). Individual sgRNA vectors were validated *in vivo* as previously described (Johnson et al. 2023) with a ubiquitous *Eef1a -1955/-1>Cas9* or *Eef1a -1955/-1>Cas9::Geminin (Stolfi et al. 2014; Johnson et al. 2023)*. Genomic DNA was isolated using QiaAMP Micro extraction kit (QIAGEN), targeted regions amplified by PCR using AccuPrime Pfx (ThermoFisher), and PCR products purified using the QiaQuick PCR Purification kit (QIAGEN) following the published protocol (Johnson et al. 2023). Samples were sequenced using commercial Illumina-based amplicon sequencing (Amplicon-EZ; Azenta) and efficiency was determined by indel % compared to controls. Data on sgRNA efficiency is provided in **Supplemental Figure 1** and sgRNA sequences are provided in the **Supplemental Sequences** file.

### Bulk RNA sequencing

Fertilized zygotes were prepared and electroporated as described above. Zygotes were electroporated with DNA plasmids using a pan-neuronal driver *Nut -1155/-1(Shimai et al. 2010; Johnson et al. 2023)*: *Nut-1>LacZ Control*, *Nut-1>Pax2/5/8.a transcript variant 1 (“tv1”,* transcript model *KY21.Chr6.690.v2.SL1-1)*, or *Nut-1>Pax2/5/8.a transcript variant 2 (“tv2”,* transcript model *KY21.Chr6.690.v2.SL1-1*). Experiment was repeated in duplicate. RNA was isolated from 10 hpf embryos using a Monarch total RNA miniprep kit (New England BioLabs Inc.) RNA was stored at −80°C until sample analysis and sequencing by the Molecular Evolution Core at Georgia Tech as previously reported(Johnson et al. 2023). Briefly, total RNA integrity levels were measured by the Agilent Bioanalyzer RNA 6000 Nano kit and all samples had RINs above 9. Enrichment for mRNAs was performed using the NEBNext Poly(A) mRNA isolation module and Illumina libraries were prepared by the NEBNext Ultra II RNA directional library preparation kit. Libraries were pooled and sequenced on the NovaSeq 6000 with an SP Flow Cell, to obtain PE100bp reads. Pax2/5/8.a transcriptional variant fold change Pearson Correlation analysis and graph generated in Prism (9.5.1).

Data quality control and analysis were performed in Galaxy (usegalaxy.org)(Community 2022). Raw reads were quality controlled with FastQC and Cutadapt. Reads were mapped to the HT_KY21 *Ciona robusta* genome with RNA STAR and checked using the Integrative Genomics Viewer (IGV; Version 2.14.1)(Satou et al. 2005; Satou et al. 2022). The number of reads per annotated genes were counted using featureCounts and DESeq2 was then used on the read counts to normalize them to the controls. Datasets were then annotated with the HT_KY21 genome (Satou et al. 2022). KY21 gene models were linked to KH gene models using the Ciona Gene Model Converter https://github.com/katarzynampiekarz/ciona_gene_model_converter. For each step, quality reports were aggregated using MultiQC. Raw FASTQ files can be found in the SRA database under BioProject accession number PRJNA981160. Analyzed data is provided in **Supplemental Table 1**.

### Single-cell RNA sequencing reanalysis

Re-processed scRNAseq data from Cao et al. 2019 were analyzed (Cao et al. 2019; Johnson et al. 2023). Data from *Ciona* embryos in the Mid-tailbud II stage, roughly 10 hours post-fertilization (hpf) were analyzed using the Seurat v3 package in R to identify cell type clusters based on unique gene expression markers (**Supplemental Figure 2A**)(Satija et al. 2015; Stuart et al. 2019). Replicates were integrated and pre-processing and clustering were performed using the SCtransform and FindMarker functions (Hafemeister and Satija 2019). Cluster 25 was identified as containing Neck cells based on high enrichment for *Pax2/5/8.a* reads (**Supplemental Figure 2B, Supplemental Table 2**). Cluster 25 cells were re-clustered based on differential gene expression, resulting in two sub-clusters (see **results**). The FeaturePlot function was used to visualize feature expression in low-dimensional space, and to use colors to map out the relative expression levels for each gene of interest in the Neck and brain clusters. Additionally, VlnPlot and RidgePlot functions were applied to the expression distributions within the clusters, allowing for the heterogeneity of the Neck cluster to be further examined and the potential for additional sub-populations noted. R objects and code can be accessed on OSF at https://osf.io/uc32x/.

### RT-PCR of Phox2 cDNA

RNA was isolated from 17 hpf *Ciona robusta* larvae by Monarch Total RNA miniprep kit (New England BioLabs Inc.). cDNA was produced by Omniscript reverse transcription kit (QIAGEN). The resulting cDNA was diluted in nuclease-free water and stored at −20°C. Based on the RNA model discovered during RNA sequence analysis, forward (5’-CATAACGATGGACTACCCTGC) and reverse primers (5’-CAGACATGTCGTGGTAGGATAGG) targeting the new exon 1 of *Phox2* and stop site of *KY21.CH14.159*, respectively, were used to perform PCR on the single strand cDNA with OneTaq 2X Master Mix (NEB, M0482S) in 50 μL reactions with a touchdown PCR program. PCR products were purified (QIAquick PCR purification kit; QIAGEN) and TOPO-cloned into dual promoter empty vector (ThermoFisher, 450640). White colonies grown on LB plates with 100 mg/mL ampicillin and coated with X-Gal were selected and tested for an insert of correct size by restriction enzyme digest. Plasmids of the appropriate size were sequenced by Sanger sequencing with M13 forward and reverse primers (Eurofins Genomics).

## Acknowledgments

We thank Shohon Rafique, Tanner Shearer, and Lindsey Cohen for technical assistance. We thank members of the lab for feedback and suggestions. We thank the Optical Microscopy Core for assistance with confocal imaging. We thank Shweta Biliya of the Molecular Evolution Core facility at Georgia Tech for processing bulk RNAseq samples. We thank Bernd Fritzsch for helpful feedback on the manuscript. This work was supported by an NIH F32 Fellowship (F32GM150234) to EDG, an ECSEL undergraduate award to LC, an ARCS Fellowship and a Mortarboard Award to SP, and NIH award R01GM143326 and NSF IOS award 1940743 to AS.

## Author contributions

Conceptualization, E.D.G and A.S.; Methodology, E.D.G., K.M.P., L.C. F.R., and A.S.; Investigation, E.D.G., A.G., L.C., S.P., H.S.A., and S.M.S.; Formal Analysis and data curation; E.D.G., K.M.P., A.G., and F.R.; Writing – Original Draft, E.D.G., K.M.P., and A.S.; Funding Acquisition, E.D.G. and A.S.; Resources and Supervision, A.S.

