## Supplemental File for "Specification and survival of post-metamorphic branchiomeric neurons in the hindbrain of a non-vertebrate chordate"

### Supplemental Figure 1

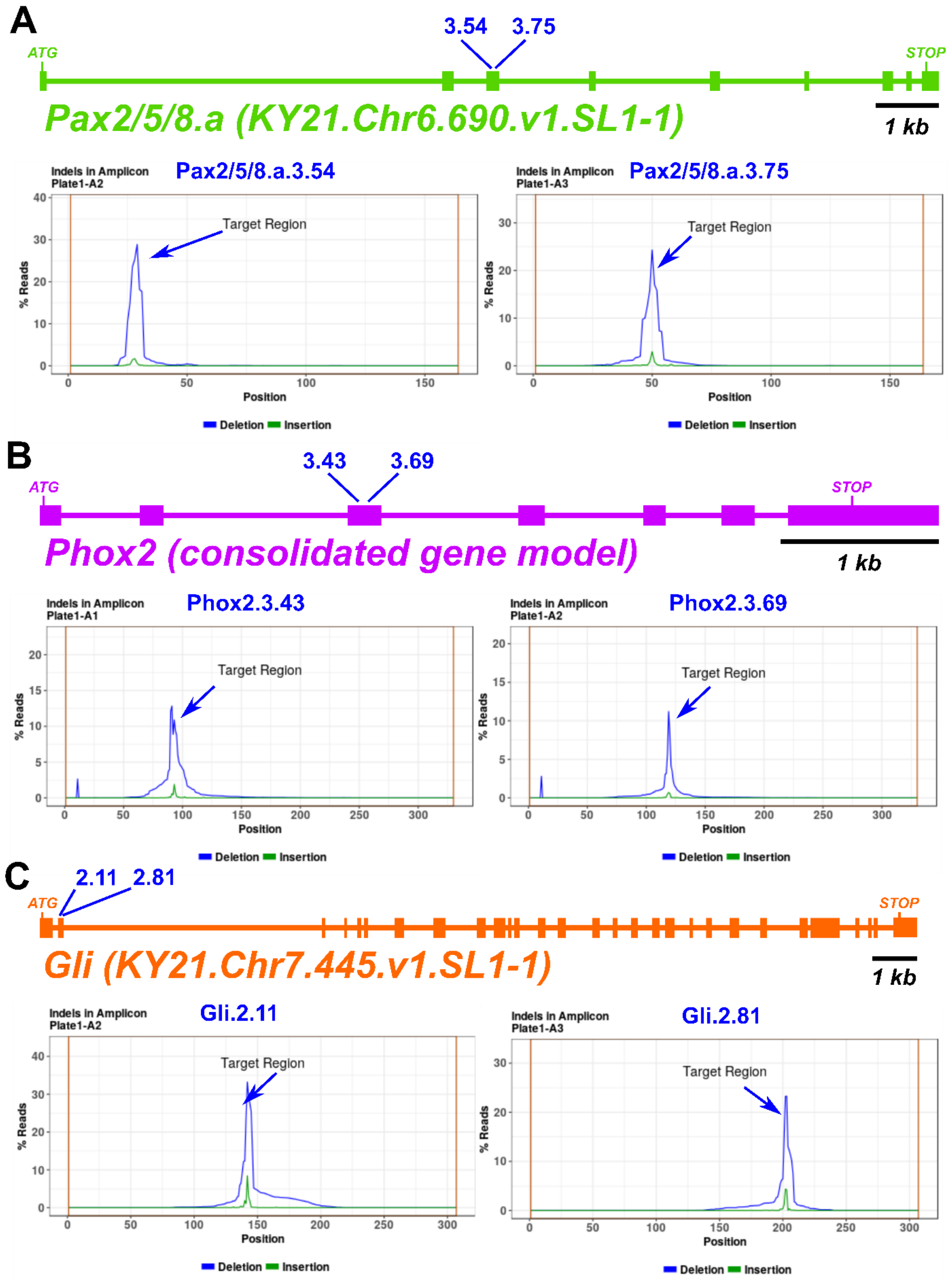


1

1

**Supplemental Figure 1.**

Indel plots generated by Amplicon-EZ Next Generation Sequencing from Azenta/Genewiz for validation of sgRNAs targeting (A) *Pax2/5/8.a*, (B) *Phox2* and, (C) *Gli*.

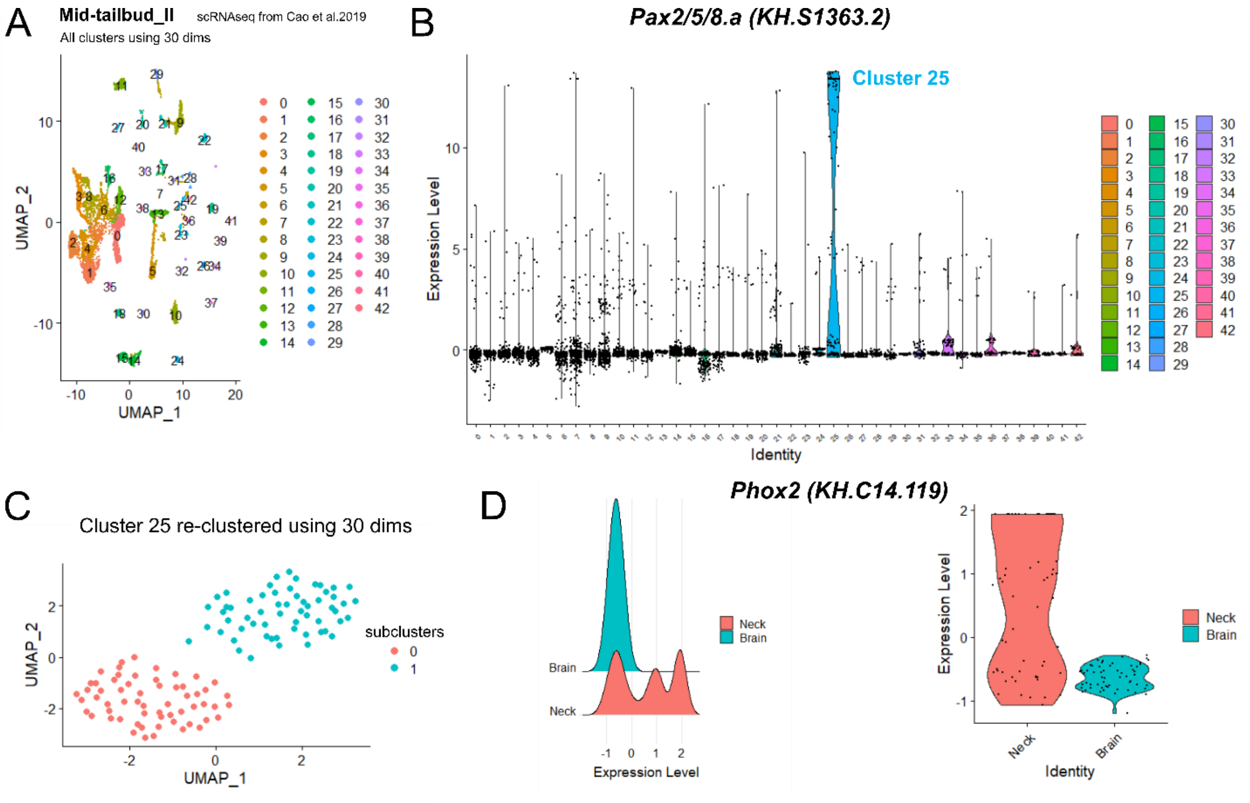


**Supplemental Figure 2.**

A) Clustering of re-analyzed scRNAseq data from Mid-tailbud II stage [1]. B) Cluster 25 shows enriched expression *Pax2/5/8.a*. C) Re-clustering of cells only in Cluster 25 revealed subcluster "0" (Neck) and subcluster "1" (brain) showing clear separation. D) Phox2 expression is further enriched in a specific subset of subcluster 0 cells, as indicated by both ridge (left) and violin (right) plots, mirroring the detection of *Phox2>GFP* reporter expression in only a subset of Neck cells during embryonic stages.

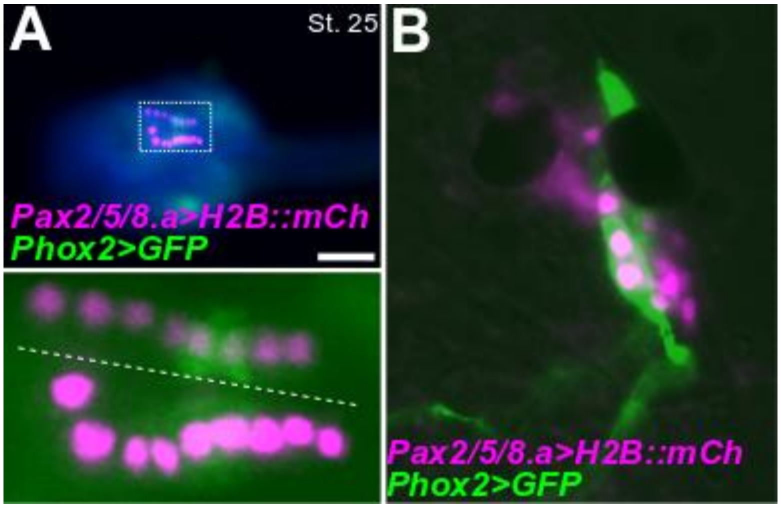


**Supplemental Figure 3.**

A) A Stage 25 (~14 hpf) embryo expressing *Pax2/5/8.a>H2B::mCherry* (magenta) and *Phox2(C.intestinalis)>Unc76::GFP* (green). The Neck on the left of the midline (dashed line) has nine Pax2/5/8.a+ nuclei compared to only eight on the right side, demonstrating that Neck cell divisions can vary between left/right sides within an individual. Scale bar is 50 μm. B) Close up of post-metamorphic Stage 38 (~96 hpf) juvenile brain shows Pax2/5/8.a>H2B:mCherry+ nuclei (magenta) Phox2(*C.robusta*)>Unc76:GFP+ neurons (green) innervating the gill slits. Individual is the same as in **Figure 2K**.

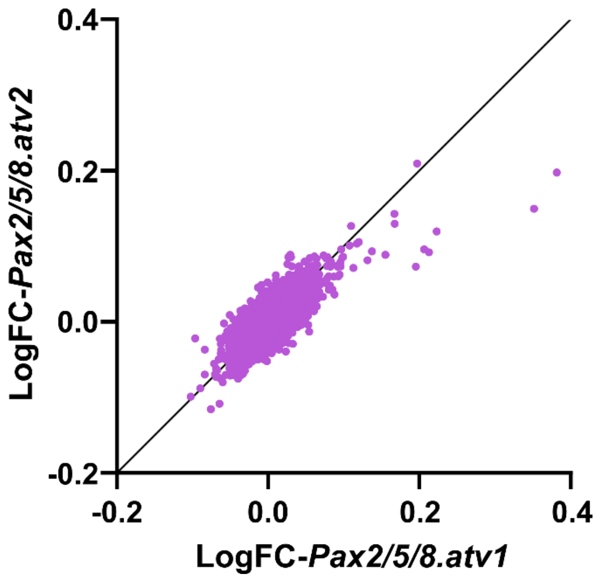


**Supplemental Figure 4.**

Dot plot matrix showing comparison between average Log2 fold-change values for all 16,252 genes upon overexpression of Pax2/5/8.a transcript variant 1 (tv1) and 2 (tv2). Raw data can be found in **Supplemental Table 1**.

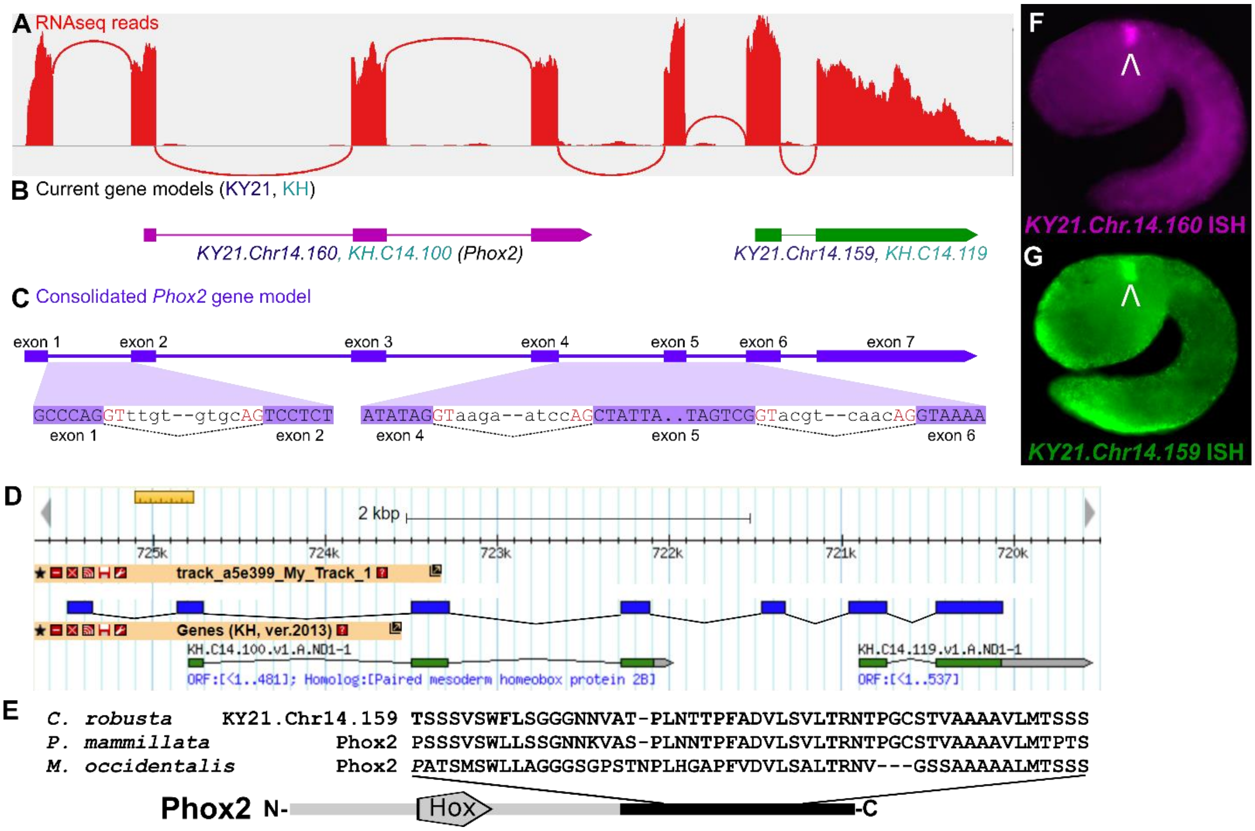


**Supplemental Figure 5. RNAseq reveals cryptic exons in an updated *Phox2* gene model**

A) "Sashimi" plot of RNASeq reads mapped to the *Phox2* locus in the *C. robusta* genome in (adapted from IGV). Red lines indicate reads spanning exon-exon junctions. B) Current Kyoto 2021 (KY21) and KyotoHoya (KH) gene models. Comparison to RNAseq reads in (A) shows that KY21.Chr14.160/KH.C14.100 and KY21.Chr14.159/KH.C14.119 should be consolidated into a single gene models, and reveals the existence of cryptic exons 1 and 5, which are not represented in any current *Ciona* gene model. C) Updated, correct *Phox2* gene model confirmed by cDNA cloning, showing splice donor (GT) and splice acceptor (AG) sequences flanking newly identified exonic and intronic sequences. D) Alignment of *Phox2* cDNA sequence (blue) to the KyotoHoya (KH) *Ciona robusta* (*intestinalis* type A) genome (green). The sequence reveals a cryptic exon 1 and 5 of Phox2, linking gene models KH.C14.100 and KH.C14.119. Panel adapted from KH genome browser [2]. E) Alignment of proposed Phox2 C-terminus amino acid sequence and Phox2 protein sequences from *Phallusia mammillata* (*Phmamm.g00000831*) and *Molgula occidentalis* (*Moocci.g00011046*)[3]. F-G) *In situ* mRNA hybridization using probes designed for *Phox2* (F) or *Chr14.159* (G) current gene model shows identical expression pattern in the Neck at stage 22 (~10 hpf), further suggesting the two gene models are likely to represent a single *Phox2* transcript.

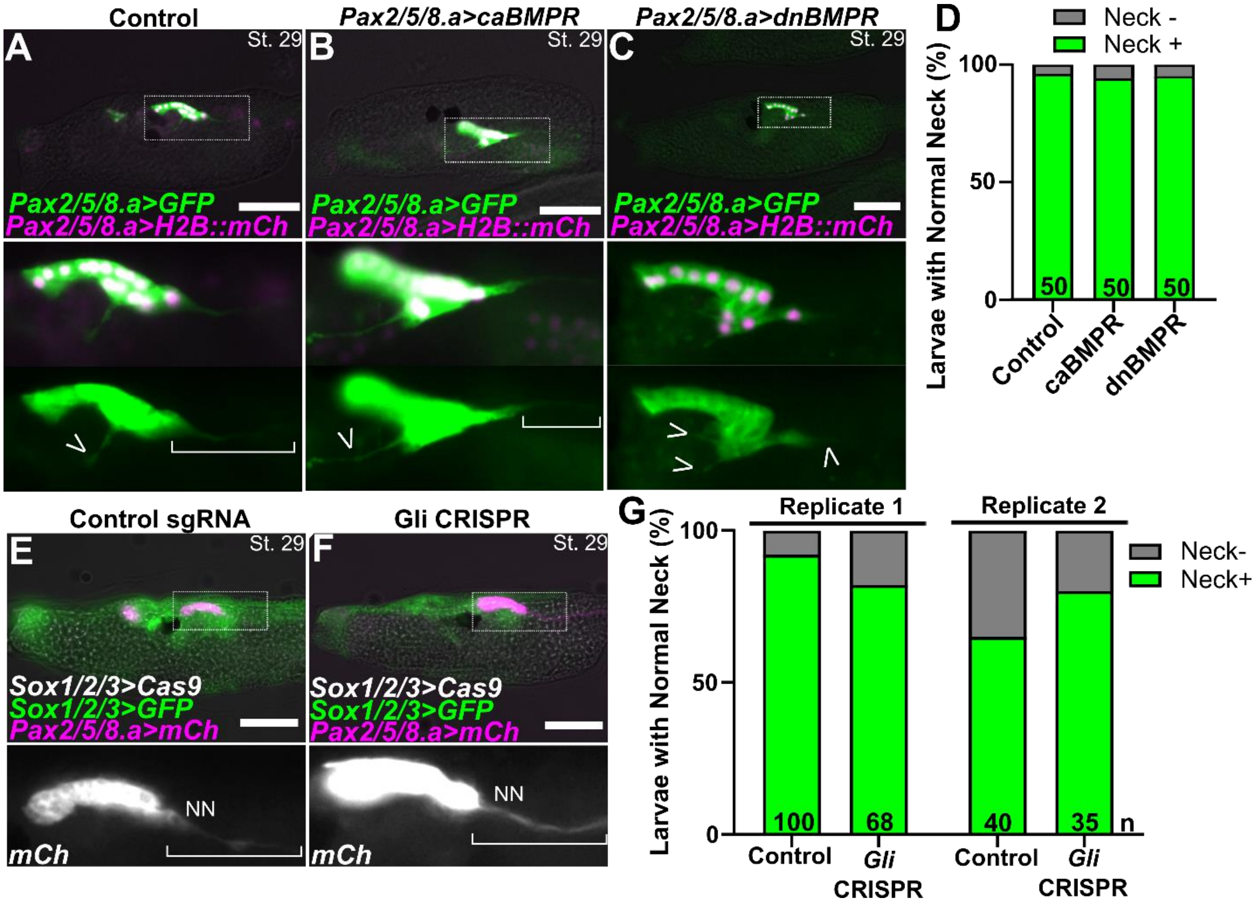


**Supplemental Figure 6. BMP and Hedgehog signaling do not affect differentiation and morphogenesis in the Neck**

A) Negative control larva electroporated with *Pax2/5/8.a>lacZ, Pax2/5/8.a>Unc76::GFP* (*Pax2/5/8.a>GFP;* green)*, and Pax2/5/8.a>H2B:mCh* (magenta) showing typical morphology of the Neck, with some neuroepithelial cells anteriorly and differentiating neurons and axons (bracket, open arrowhead) in the posterior. B) Larva expressing a constitutively-active BMP receptor (caBMPR), showing no noticeable difference in Neck-derived axons (bracket and open arrowhead). C) Larva expressing a dominant-negative BMP receptor (dnBMPR), also showing no noticeable effect on Neck morphogenesis or patterning. D) Scoring for "typical" Neck morphology in larvae represented in panels A-C. E) Negative CRISPR control larva expressing *Sox1/2/3>Cas9:Geminin*$^{N\text{-}ter}$ and *Sox1/2/3>Unc76::GFP* (*Sox1/2/3>GFP*, green) reporter, with *Pax2/5/8.a>Unc-76::mCherry* (*Pax2/5/8.a>mCh*; magenta) labeling the Neck including the axon of the Neck neuron. F) Knocking out the Hedgehog effector-encoding gene *Gli* also does not alter Neck morphogenesis. G) Scoring of larvae represented in E and F. In panels D and G, n = number of individuals scored in each sample. All scale bars = 50 µm.

**Supplemental Table 1. Differential gene expression in Pax2/5/8.a overexpression conditions as measured by bulk RNAseq.**

**Supplemental Table 2. Transcripts enriched or depleted in Cluster 25 of mid-tailbud II scRNAseq re-analysis.**

**Supplemental Table 3. Transcripts enriched or depleted in subcluster 0 (Neck) vs. subcluster 1 (larval brain) within Cluster 25 of mid-tailbud II scRNAseq re-analysis.**

**Supplemental Table 4. Cross-referencing subcluster 0 scRNAseq differential gene expression and Log2 fold-change from Pax2/5/8.a overexpression bulk RNAseq.**

**Supplemental Video 1. 3D projection of CMNs in post-metamorphic juvenile shown in Figure 2L.**

**Supplemental Sequences**

Some sequences are based on genome assemblies and might not be verified by sequencing after cloning. ANISEED unique gene IDs of marker/reporter genes are at the end of the document.

>Pax2/5/8.a -1961/-1 [smaller version of reporter first published by Oonuma et al. 2021[4]]
TGTCAGCTAGTGCACTCTTACAGTGTTGCCAGCTTTTGTTTTTGCTGGGCCTAATGTTGCCAGCTATGTC
CCCAAGCACTTCTCAGTGCTTCTACATGTCTTTGAAACAAATGCAAAGGAATTGTTGCAACAATCTTACT
TGATAAAGATTAGCATCTACTGTAATGTTCATTCCTTCCGAGTGTTTTGTCTGTCGGGACTTGAATCTTA
CAGCGGTTTTGTAGTGTCTCGGTTTGCTTTGCGAGAACGCTGTCCCAGTATACATATACTCGAGTAATAT
CAATATTTAAATTCCTTTAATTCTTTAATAAATTAAATTAGTATAAAGTCCCCAACACACCGCTAAAGTT
AAATTATTTAACACAACATCGCGTTTGGCATAATTGTTACGATGGCCACGTCACAATTACGCTTTACGTC
GATGCGGTGGGTCAGCTACGTCATGACCCTGCGTTCGGTGCGGTTACGACACGGTCGGCTTAGTTACGTT
ACAGTTGCGGGTTTGTTACGTGGACGTCGCCGCTTTGCGGTTCGTAACAGCCACAGTATGTCGTAACGGT
CGGTGCGGTGCGTAACAGCCGGACGATATTATATTGTGCGAAGATACGACGAGAGTATCGCGGTACCAAG
ATGTTTTCTCAATAAGCGGTCGGGTGATGCAGGGCCGTTGTTTGCTTTAGACTTCTTGCAGGTGGGCACT
TGGTGGCGTGCGCACTGCAGGGCTTTTCTGTGAGACTTGCAACTTTTCGTTTGACGGTGAATGCTTGTTG
GTGTGACTGTTTTTGTCTATGGGTTTCTTTTCCTTGCTTGGGTAGGGTCTATCCAGCAAAGGGTTTCTCA
CCAGAGTAGAAACCCACTACCATTGGCTGGGGGCTTGTCGTGCCGTAGTGGCTTTCCACCAACGCCAGCC
ACATAAACTATGAAGGTATTTTAATCCTGTTTTTATTTTCCAGAATTTCAGTTTCAGATATTTTCATCCG
CACGTGGGCGCTAGCGATTGTTGGAATTACAGTTCGCATTGTTTGTAATTGTTATAATTGTAAGTCGCTT
TGTGCTGTGACGTCATAATAAAATATATTTCTGAACTTTGATTACGTCATATTCGTGTTGCTTAATGATG
AACTCGCGATACTAGTGACGTAATAAATGAAATTTGATTTTAATTTGATTGTTCTGTAGACGCCGCCACG
CCGAAGTCCAACTGCGTCGAACTGACAATACGCTGTGTGACGTCATAATGAAACAAGTGCATGAGGTAAG
TGACGTCACAATCGTGGATTAAGAGCGGAAGTTATCGCGTTACGTATTATGACGTGTACAATGTTTATTG
TTTCATAGTTAATCGGACGGAACAAATTAACTTGTGACGTCAGAGGGGTCAAGTCGCTTTATTTGTTTAA
ATTTTAGAATTGTTTAGCTTTTGAGACTAAACTATTGTGACGTCATAAACGAATTTCGCGTCTTCGTTTG
TCAACGTTTTTGTTACTTGTGACGAAATAATAACTAATTAGAAATGCTGACGTCCTAATGTATTACGTAA
TAATATTGTGATGACGTAGCATGATAATGACGTAGCTCGGGGTAAAATCTACGCCTAAATTTTAAGAGAG
AGCTTAAAAAAAACGAGAGCGGCGATAGAAAAAGGCGACTCCCTTTTTTCCACCCTGTCTCTTATATGTT
GGAGTGAAATCTTCGCATCTTCCATAAAGTTAAAAGTTTCGAAGATTTCACTGTTGGGTGATCGTTGCAG
AATAATCACTATAACTAGGGTTTGTTACGTCATAAAAGATTTAGGGATAATTATTTATTTTGTAGAATTC
TTATGCATTGTTACGTAGTAAGCGTTACGACGTCATACAGCAACCGCATTTTGTCTATTTATCGCAAAAT
TAGCTTTCTGACGTCACTGCATTATGACATCATCATGTATTACGTCACACAGGTGTTAGTAAGTTTGgAT

>C. intestinalis (Type B) Phox2 Promoter -2951 to -1 [Dufour et al. 2006[5]]
CCCATCGGCGTTTTAAAGTCGGCTATTCTCTCTCTCAAATTGGAATTAACTCTAAATTGAAGTTTATGCG
GTATCCATATGCGAAATATAGGGCTCGTTTATTGGCAGCATCTGGCCGAGTGTTTCGACCACCGGCAATA
ATAACAGCACACCTACGCTCCACTGTTAGAACGTGAACCGGTCTTTGATGGCCAGAATAACAGCTCGGCT
TCGCTAAACGACATTTGGTGAACGATGATACTTCTTAGGTTTAACCGAACAGCGGGGGGCAGCTATACCC
GAACGCGACTCGTATTCATTTTCAGCGAGTTATTGTATCATATAGTAGGGTGGGGGAAGATGGGACACCT
TTAGCACATAATGTTTAAATATCCTGATCGTGTTTTAAACAAAACAATGGTCTATGGGAGTCGTGAGAAT
GCCGTTATATAATTCGTTGAATATTGTTTGTTTACTACCATATGCAACGAGAAAATGGAATGAAAAGGTG
TTCCATCGTCCCCCACCCTACTACACTGAATGCTTTACCGACAATGAATATTGGTTTGGTCATAGCACGC
TATAACTTCACGCTATGTAAAGAAGTGTGGATGCGAAAATTCTCATGATGTAAATTATAACAGACAACTA
ACTAATAGAAATGAACCCGTTTCAACTTTACCGCTGAGATATTCAATGAGGCGTTATACCGATTCCGGAA

TTAGTTGATAATTGCTTGAAGTTCCGAATGCGACACCCAAACAAAGGCACCCCGTATACCCATTTAGTGT
TACAAGCCTTGTCAAACAGATTAATGAATTTAATTCGCATGAGAAAATAATGTTTATTACATTTGAATAT
TGTTTCCAACATTTGAAAGTTAAAAATTCCTTAAAATCTAATTGGTTTGAAAATCCTTATCAATTTTTAG
AGAAATTTATTTTTCATTTAAACGCTTCCTAGTAAAACAAAACCTAGGCCTAACGTGGTTGCTAAAACAA
AATGGAGACGGTCGGTGCTCGTACAGCAAGAGAGCGAGCCAATAACGCACTCACGAGAAGCAATTAGGCG
CGGAATCAACTTAATCCAAGGTTTCTTCGATAAAAAGCCAACACTGTGTTGTACGTGAGTGCTAACATCG
AGTCAATTCTAACCTGGTTATGTTTGGTGCGCTGACGTCCTCGAGTCGGTAGAAATGTCATTAGAATTCG
ATTCGAAATGGAGTTTATTAAGTAAAGGCGGCGGCACTCGTCGGCCAGCACCTGGCGCTTTTAGTACTGG
AAGGACTGAACAATATACTGAGAAATAGCTTCCACACTTCCACATTTTACAAAAAAAAAATAGTTATGGA
ACACATGTGTTATGTTTTAAATAACATAATAAAACTCTTTGAGTAACGAATTCAGCGGTTTTTCAAAACA
TTTTTATCAAAATTATGTTTTTCTTATTAAACAGTATATTCTTATGGTTTTGGAAGATCTATATACATTA
AAAACACGTAGGATATATTACGGTATTTTTAACATTTTAAATAGTTTTATGTAGGTTTCTTGGTAGAATG
CGATGATTTAAATTCAATCCAATGTATTTCGAGATAAATGCGTTAACAATTGTGTGTTAAAGAAAGAATC
TGCAAACATATTTTGCGGTACCGTCGCTTTTGCCGGCCGGATTTAATCCAATTTGATCATAACGCGTGCC
ATGGTTCCGACCCGAGGCGAATCACTGGTGGGGAAAAAGGGGAAAGGCCAATTGACTTTATATCGGGAAA
GCTAACGTTATTGGGCGGACAGAGCTAAAGCGTCGAGTCATTGCTGTGGCCTAATGCAGTTTAGGTGCGA
AATTGGATTTTGCGATTATGCAGTTATGATCGGAAAGTTACAGACAATCTGTCTCAGCACATATATTAGA
TAACTTAGATTTTCAACAAATTTTCGCTCATGCTGTTTCGGTAGATCGGCTAACGTAATAAACAAATAAT
TTCGTTTTCCGAAAGCAGCATCTCTACTGCATTTGCCTGCCTAACAAGGATTAGATTATAGAACCTGGAA
AGCGACAGTGGCTTAAACGCCGGAATAAAGTGATTAATAAAGCAAAACAAATTAAGTTCAAAACTTAAGG
GCGAATTGCCTGAGGCAGCAGTCATTTTTTTAAAGATAATTTTTACATTTTTGCTATAAAACCTAACCAT
ATAAAAAATTTAAAAAAATCAAATAATAAAACCTTTTATTTTACAGTTGAAAAAATTAAAATATAACGAT
ATGGATTACCCTGCGTATTTAGGGGTTGGCACAAACTATGACACGACTGCTTGCATGGTGGCTGCGGCAG
CTAATAATGATCCACACCAACCATACGTTACTACACCATACGGGGATTTTAACTCATGCGCTCACGCCCA
GGTTTGTGCTGCATTATTTTACCACGCATGGTCTAAAAAGATATTAATTTCTGATAAAGTCGAAATATAT
TTGGGCTTGTAAAATTGTTTTATAGCGCTGTGGGGTAGGATGGGAAACCTTTAGCACCTAAAATCCGTAT
TTCTTGTTCCTGTTTTGAACAATTAACAACGATCTTTTAGAGCCGCAGGGCTACGGTTGTATAATTCTAT
AAATATTCTTTGTTTAGTATGCCAAATGGAACGAGAAAAGAGAATAAAAACACGTCCCATCTTACCCCAC
CCTACTATAACTTTAATTTCTGGTTAAGTCTAAAGAGATTTGAACTTATAACATTCGTTTTATTGTATTT
TTCAATTTCAATTTAACATTTTCATTCATTCGCATCTGCGCCTATTCTTATAATTCAAACAATAATGTCT
GCAAGTGGTGTTGAGCTTACAGTAATTTAAATTCCGAGTTATTTAAATTGAAAGTCAATAACCTTATGTA
CAGTCCTCTCAGAGCAACTTCGCGTCAGCTTATACACCGACAGTCCCGATCCGGAACTCCAGTTTTGGTG
GCGG

>Ciona robusta (Type A) Phox2 promoter -2416 to +15, coordinate based on newly identified start site in exon 1. ATG start codon

CAGAACGACTTGCTCTGACAACTTAATTATAAGCGGCTGGTCAGTCGCATGCGCGGTCGACCGATTACCA
TTTCCGCCCGCGCAAAAATCAACAATTTCTTGAAAATAATAAGAAAAACTTTCGGTGAATTAGCCCTTTT
CTATATACGATATAGACACATTAGTATTTTAAAGACGGTTATTCTTTCTCACAAATTGGAATTAACTCTA
ATTGAAGTTTATGCGGTATCCATATGCGAAATATATGGCCCGTTTATTGGTAGCATCTGGCCGGGTGTTT
CGACCATCGAAAATAATAACAGCACACCTACGCTCCACTGTTAAAACGTAAACCGGTCTTTGATGGCCAG
AATAACAGCTCGGCTTCGCTAAACGACATTTGGTGAACGATGATACTTCTTAGGTTTAACCGAACAGCGG
GGGGCGGCCATACCCGAACACGTCTCGTATTCATTTTCAGCGAAATATTGTATCATATAGTAGGGTGGGG
CGATATGGGACACCTTTAGCACGTAATGTTTAAATATCCTGATCGTGTTTTAAGCAGTTAGCAACATTCT
ATGGAAGTCGTGAGAATATAGATCTAAATTTCTTTAAATGTTCTTTGTTTACTACTAAATGGGATAAGAA
AATAGAATGAAAAGGTGTCCCATCTTCCCCCATCCTACTACACTAAATGCCCAACCGAATACACAATAAA
TATACCGGTTTGGTCATAGTATTTTAACTACACGTTATATAATGATATGTGGGTGAGAAAATTCTCATGG
TGTTAAATTATAACAGACAACCAACTAATGGAAATAAACTCATTTCAACTTAAACGCTGAGATATTCAAT

GAGGCGTTATAACGATTCCGGAATTAGATGATAATTGCTTGAAGTTCCGAATGCGACACCCAAACAAAGG
AACCCCATGGCCGTTTAGTGTAACCAGCCTGGCGAACACATATTATGATTAATGAACTTAATTCGCATGA
GAAAATAACGTTTGTTATATTTAAATTCTAATGCTAGCATTTAAAAGCAAAAATTTCACCAAAATCTGAT
TGGTTTGAAAATATTTTATTAATTTTTTGGGAAAAATATGTTTTATTTCAACGCTTCTAGTGATACAAAA
CCTAGGCTTTAAGTGGTTGCTGAAACAAAATGGAGACGGTCGCTGCTCGTACAGTTAAGCGAGGGAGCCA
ATAACGCACTCACGAGAAGCAATTAGGCGCGGAATCAACTTAATCCAAGGTTTCCTCGATAAAAAGCCAA
CACTGTGTTATACGTGTGTGCTAACATTGAGTCAATTCTAACCTAGTTATGTTCGGTGCGCTGACGTTCC
CGAGTCGGTAGAAATGTCATTAGAATTCGATTCGAAATGGAGTTTATTAAGTAAAGGCGGCGGCACTCGT
CGGCCAGCACCTGGCGCTTTTAGTTCTGGGAGGACTAAACAAAAAACTGAGAAACAGTTTCCACACTTTT
AAATTTTAGAACAAAAAATATATGTTTATGCAACATTTTGTAATGTTTTAAATAAATTAACAAAACTCTT
TAAGTAACGAAATCGACGGTTTTAAACTATATTAATCAAAACTAACTAAGTATTTCTAAACAAAAATAAT
TCCTTATGCTTTGGAGTATCTTCATACATTCAAAAAAACGTAGGCTATATAACGGTGTTTTTAACATTTT
GAGTAGCTTTATAGATTTTTGGTATAAAAAGCGATGATTTAAATTAAATCCAATACATTTCGAGATATAA
ATGTGTAAACGTTTATGTGTTAAAGAAAGAATCTGCAAACATATTTTGCGGTACCGTTGCTTTTGCCGGC
CGGATTTTAATCCAATTTGATCATAACGCGTGCCAGGGTTCCGACCCGAGGCGAATCATTGGTGGGGAAA
AAGGGGAAAGGCCAATTGACTTTATATCGGGAAAGCTAACGTTATTGGGAAGACAGAGCTAAAGCGTCGA
GTCATTGCTGTGGCCTAATGCAGTTTAGGTGCGAAATTGGATTTTGCGATTATGCAGTTATGATCGGAAA
GTTACAGACAATCTGTCTCAGCACTTATATTAGATAAGTTAGATTCTCAACAAATTTTCGCTCATGCTGT
TTCGATATATCGGCTAACGTAATAAACAAATAATTTCGTTTTCCGAAAGCAGCGTCTCTATTGCATTTGT
ATGGCTAACAAGGATTAGATTATAGAACCTGGAAAGCGACAGTAGTGGCCTAAACGCCGAAATAAAGGGA
TTATTAAAGCAAAACAAATTAAGTTCAAAACTTAAGGGAGAATTGCCCCAGGCAGAAATCATTTTTTAAA
GATAATTTTTAACACTATTTGCTAACCAACCTATCCATGTAAAGAAAATTATAAAAATCAAAATAAAACT
TTTTTTTACAGTTGAAAAAAAAATAAAACATAACGATGGACTACCCTGCGTAT

>Sox1/2/3 -2391 to -1 [Stolfi et al. 2014[6]]
CGCTAGCATGTCAAATAGTTCGAATTTTATTGTAGAACTCGGGCAGTTGTGATGTAACACGTGAATTGGA
TTCAGCATTCGTTCTTTAAACCTATGAGTCGGCAAGGAAGCGATCTGTCAAATTAAATCGAATCTTTCAG
CGCTCGCATCGAGACATAGCGATACTGGAAGCGTCGGAAATAGAATAGTGGGGTTGATGACAGGTGAAAA
CAGAATTCGTTGCGCAAATTACGAAATATTTTGCGCTGGTTTTATTACACAACAATTTTCTCAATAGTGG
GGTCGAAATTGATGTTTGCATTTTGTTGTAGGGGTTGTACATGCGGGATACGCTCCGTGGTAGGCGTCTA
TGTTATACAGTCGGTGTAACTTAACGACAAATATAGAATACGTTGTGTTCGTTCATTGATGGTCTTGAGT
TGCACGATTTCCCGTCGCAAATGAACCGTTAACACAACTTGGCCCGAAGCAGGCATGGCAGAAGTCGTCG
TAAGTAGGCTGGGGTGTCGTGGGGCCGCGCGTCGTCATGCTAAAACCGACCGAGCAGAGCGTAGGCAAAA
GTAAGCACTGAATTGAGTGAAACGCGACGAAAAAAACGGTGGTACTTGTCGGTTTGTCCGCGCTGAAACC
TAATAACTGGCTGCGCGTGTTTTCTAGGATGTCGAGTGTTCGCGGACACTTGAACACTTAGATATCTACA
TGCTGTCGTTGCTTTGGTTGGTTTTACACTGGATTTGTATTGCTGGTGTTTTGCTGTAAAGCCAGAGTTT
GTAACCATATAGACATCGTATATTTGTAATGTAGTATTCCTAGTTTTGAGTATAGAGCTCGCTATTTAGT
TCTGTTGTCAGTTTTAAGTGTGTTAAGGCAAGATACATAAAGAAACAAAGTCTTTATCTATCTTGTCGAA
TAGAGTAACGAAGATGAAATTATTAAAAGTTTAAAAACAAGGAATTAACAGCAAAATGACAGTTGGTAAA
ACACGCTATGAATAAACTGTATTGAAAAACATCAGGAAATCGTTTGGTTTGTGTTCAATTTGTTTAATTT
AGAAGTTTTGTATTGTTAGTATAATGACTGTTGTCTGTACACAACACGTGTTGACCACTATAGTAGTGTA
GTAGCCCAATAACAATTACTGATGTTCTGCGCATAAGTCGCGTGTGCTTTACTGGGCATTCATGCCGGTA
GAATTAACCATTCATCGTAAAAGAACAAAACGGTCTATTCTGATCCGTTCATTGAGTGGCTTATGAATTG
GCAAAAGTGCTTGTAAAGTCGGCACTGAATGGGCGCTGCCTTTGTTACTGACAATGTGCATTCATGACGA
TAGTTCCGAAAGGTGTGGAAGTAAAATACACTGTTTGTGTTCATACTACGCATTCCGAGGAGGAATACCT
TTTTGATACGCCATGTATGCTTGATTTTTTTTGACAAAATTAAACATTACAATTATTGGAACGACCACAA
GGTTGTTAGTTTTACACTTTGTTGATATTTTTGTATGGGTGTTATACGTTAATGGGGATGTTATTGAAAT
ATAGAGTTGTGTAATTGTAATTGTGCGGTTTCATGTTAACGCTTAAAACAGTTTTCTTTCTAAGCTGGTG

TTTTGAGTTTTAGGTATATATGCTAAAGGTTTAAGTATTGGGTTATGAGTTTTGTTATTCGTTTCTTTTT
TGTAAATATAACTTTTGCGGGTTTTCATATTTTTCGATTCGTATAAAATCGAATTCTGTCTTATCATGAA
CCGACCACGTTTTTCGCTGATTCGAAGTCGTTTAGATGGTTGCTTTAGGGAACGCTGGATCCCAACGAAA
AGAAAAAGGCGACGTTCGCCTCCGCGAAGATAAGCCGTAGCGAGAGAAAGGTCGTTTGTAGCGCAATAAA
GGCAGCCTGTGAGATGCCTACTTCATTCATTCTGCCTTTTGTGACTTCATAGTGGCATTGTGAGCTTCTA
TTCAGGCATTGTCTCCAAGAAAGTTATGAGTCCATAGATTATAAGACCTCTTCTATGGACAAGCAGAACA
ATTGAATTACAATTGTTAAAAACAATCGTAAATTGTAGACCGTAGTTTCAATAGAATTAACGCGAGAATA
TTTCTGGAGTAAAAAGTCTGAAATAGAAAAATAGGGACCATTAAAAATTGTCAGCCGATAGCCAGAATAT
GTTACAAGTCAACTGTGATTTGTGTTTGAAGTTTTTTAAATTGATTTTTAACTGAAACTTCTACTTTACA
GCGTTTGGAATCAAGTAAAGATATTTAATTCAATTCTTGCGAACATTCGCCTAAAGTCTCACACGTCATT
AAACTGGATTTTGTAGCTTACAAAAACTTCTCCGTCCCTACTCCACCGGGGTTTCTGAAAGAGCCATCTC
AGAACGACTTC

>Nut Promoter -1155 to -1 [Shimai et al. 2010[7]]
ATCTGTTCTAGGAATCTTTTAACTCGGCGAGCCTGTGTTTTCTTTTTAGACTATAATACCCTCCAATAAC
AGGGTAAGATGCATGCGAAAAAAACGATTGATTTTGTGCATATAAAATATTTATAATAGTCTGGGGTTAT
GTGAGTGAAATAGCTGGTTATATCTAACTGCAGCCGGAAATACGTGCTCTTTGCAGAAAGGAAAATCCAT
TTCTCGTGTATTGCAACATACGGTATAGTAGGTTTAGATGGGTCACCTTTACCACAAATAACATCCAAAT
ATCCTGAGCATGTTTTAAACATTTAACAACGGTATATGTGGAAGTTGCGAGAATACGGTTTGATAATTCA
TTAAATATGCTTTGTTTACTACAAAATGGGACGGGAAAATAAAATAAAAAGGTGTCCTATCTTCCCCCAT
CCTACTATATATATCATATTTTCTAATCCTGTTTTAGCAATTAACAACGCTCGTTTAAAGTCGTGGGTAT
ATGCGGTTACATAAATTTAAAAATATTATTTGTTTACTTCAAAATAGAACGATAAAAGCGTGTATAAACA
TGTTGTATCATCTTACCCCATCCTGCTACAAATATAAACAATTTCATGATAAGAAAATGCACATGTTTGT
TATGCCGCGTGTTGTGTACAGTTCTGGTTAGGATTTCAGAACACACACACAGAAAATCGGTCGTGCCTGC
TGCTGCAACTAGTGCGACACCTCAATAATTATGAGACTGCCTCAGTGCATTCCTATCAACGTGTTTTGCG
CTCGTATCACCCAAAAAGCGTGCCTCGTGCTGCCTACACGGTGTGGCTACATAAAAACGTCGCGTGTTTT
ACTGTTCCAGTTCGTTTTTGTATTCGTGGAAATAGGTTCGCATTTTTTATTAAGAAAGAGTTTGGTTTTC
GAGGTCACTGAATTGCAATTAGAAGGCATTTAATAGTAGCACTAGGACACTCTATTCACTGCGTTGTCTA
AATCCATGCGAAACAACAAAATAAGTCGCAAGACATGCTGTGCGTGTATTGTTTTTTTGAAACCCCGTCG
CTTTGTGAAAATCTGGTTGATTATTTTTTCGTACCAGTTTTACAGTTTAAATACGACTGTGCTTCAGTTT
TTGTTAGTATTGAGTTGTACACTATATCAACAAC

>Gli -3339 to +57 atg start codon
GGTGATGACTGGAACCGATGACCCGTGACCGTGCGCCGTTATGCTCCGTTGCTCGCGTCAGTCGACAGCT
AATGCGTTAAGCAGTTAGCCGAAATCCTTTGCACACTAAGTCTGCTGGAGATTCGATCGAAAATAATTTG
AATATTTTCACAGCCCTTGCTTGGCTGATATTATATTACCAACCAGCAAAGCATCCAGTTTATCCTAGCG
AGTACCACCGCCGCAGGCCGTATTTGGAATCAACGTGAACTTGATACGTTTGCTAAGATATCGTAATATT
ATCAAAACTGGGAATAATGCTAGGATATATATAATCTAAATATCTTCGCTTACCATTTTTGGAAAACTTA
CGGTAATGTCATGTGTTGTACAAGAAATGTCGAAAGCGTTTTGGTATTTGGTAAATTAAACAAGGTGAAG
TGGGATAATTATAACACTTAACATTTGAAAATAACTTTCCCTCTTAAATACCACCTTGATCATAATAGTT
CGACAATTCATTTGAAGTCGAAACTCGCTGTAATTTACAACAGGTTTCCGTAATGAACAAGTGATCATTT
TTGAAGAGGAGGCGTGAAGTGAAGCTTCTCGAACTTCGACCTCAACAAAAAGTGCAACCGCCCAAAACGG
CCGGTATGCTGCCCCTTGCGGTCTCATATGTGGGCATTTGAAGGCCTAACACAGGCCTGCAATCAAAGAA
ACTACGCTTGCGTTTAGGAGATTTTCGGGCGCTGTCTTTTCCAGTTCCGCAGATTTTTTTCAGACAAAGC
GCGCTGGCCGTCACGCCTCACCGCACCCGTGTTACCATTGTAGCGTCACAAACCGAAGTTTTTTCGACGC
GCTCTGTTTATCGTTTATTATTTCGTCACGAAGCTGGCGCGGGAGCCTCCGATGCACTGCGCAAACGATC
TGTAGTAGTTAGCTATAAAACAAGCTTTATTTATATCTGTATTAAGGCTTTTCAATTTACAAGGCACATT
TTGTCGAACCAGTGTATTATATTAATCATTATTTGATAAGAAATTCGCCGAAAATGAAGAAGTTTTAGTT
CCAGTTGCAGAAAAGGTGAGTTATCACAATGCGGGTCTAAAACTAAGTGTTAGGCTACTATAATCAGCTG

TTGAGGAATGCAGTTTACGATTCCGATAGAAAAACATGTTCAAATACAGAGCTAGCAGTGTGTATAGGTA
TGTAGAGGTGCGTGTATGATGTATATTGATGCGGTAGTCTGGTGGAATGAGGTATGTAGGCCTCCCATCG
AAAGAAACGTTTTATTATTCAAAGACTGTTCGATCAAAGTCGAGGTAGAGGCGCCTTCGTGAATTGGATT
GAAATGAACCAGAACCAGCAGCAGCGGAATATTTCAGGCTCCCCCGGGGCCCGTTAATGGCCCTGCGGCT
TGTGCTCTGAGGTAACTCTGTTATTCCTACGCCTAAGCTAACTTTATAAGAGTATATGCGATTTGGGGTA
GTTCGTCCGTTTTATGCGCCATTACGACTGTTGGTAGACACCGTACGGCTGTGGCTGGTTTGTTGTTCGT
CCTCTAACTGCTCATTATGTCTTCCCGAGTGCCCGAGCCTCAATTACAGGATTACTCGCGAGTTTTAAAA
GCTTATCACTGGCGCACATAAGCCACCATCGTCACAGTACAGCGATTTCGACAAAAATGCCTGAGTGCTG
CCTCGTGGGCGCCCCTCTCTGCCCTATAACCGATCAAAACAGCATGACTCGTGTTGTATTTTGACATTGT
CGAGGGAGTTTTTCTGACTTTGTTGAAGTTTACGCTAAACGAGCTGAAGCAACCTTTCGCCTTTCTAACA
TAATTGCAAGCGCCCCGTAAAAAAAGTATTTTTGATTCGGAAACTCAAGGGCGAAGTGTTTTCAGGTTAC
GTTGGAATTTCCTGCTCAACGGTCTTGTAGCGAAACGTTAAGAGCCATTCTGCCTTTAAAATGTAAACCA
TGTATCCATTCAACCGGCGAGACACTGATACACTGATAGTCTATGCGGATTCCCACAAACAAACTCGGTT
CCCATAATGCCCAATGTATCGAATTCAAGTTCAGTAAGTGGCTTGTATTAAGTCGCATTGTGGTAGGGTA
TTAAGTTACAAGCCAAGTTCAAATACTTAAACCAGGACCACATCTAAGCGCATTTGACCGAGATCGTATC
AACTTTGGTTGATCAACGGAAATCGGGCCAATTCAAGTCCGTTAAGAATCCTCGCTTCAATAAATCTGTT
ATGTAACGTTTCCGTATCAAAAATCTTTCAGGATATATCGCTAACGGCCTCCTTCACCCCTATGGCTAAC
TATTTTTGGCTAGTAACGGTATTTTTAATAAAGTCTCTCGCCCCAATAGAAATGTTGACCCCTATCTGCC
CCACTTATAAAAGCGACGATCGCTGGGTAATCAAATAAGTGGAGTCAAGATCGGGGTCAAGATTGGGGTC
AAAAGGGTCAAGGTCAGGCGTGACCTTTTGGCTAGCTGGTGACACAATCAAGCCAAATGTGACAACAATT
GAGAATTTAATTAATAGAAATTTAGGATCGATGGTTGATTACGTCATACATCATCATACCTCGTGACGTC
ATAATTACGTCAGCTTAGATAGATTTGTAAAAGCTGGTTAGGTAAAGTTTTCCAAGCAAGAAGATACGAA
AATTATAAAATTGTTTAGTCAATCAAAATAGAGGTGTCAATTATAGCAATTACAATTTTCTGGTTACAAT
TTTTGGCATTTCCTTGAACCTATTTTCATATAGTGCCGTTTTAATCTTCGTTGGAAGTATTGAACATATT
AAAACTAATTAAATGCGGCATTTCTTGGCAGCGCTTGCCCTTGTAAGACAACCCGCAAGATCGTTCTATT
ATCTGGATGATGCCAATAATTTCTGGGTCGTTTCTCTCGAATTTCTTCCCAGAATGCGACGTCGGGCGTT
GATGGAATGTAACCTATTCTTGTAGACCGCCATTAATTTGACACCTACTGTTGCCAGGGCCTTGTCCCGT
AGACTTGCGGATTTTGCAGCTTTTGACACAGATTTGACTGTGGGAATCTTGCATGCCTTTAAACGTGGTT
TAGTTGCTCAATGTACCACAGCCGCGAATGAGTCAGTATCGAAGCCTGGCGAATGGAATTTGAGTTCTAT
CCATGTGGCGTGCCGCATCGCAAATCCGCGGCGTCTGGTTTTCGTCCCTGTTATCTGTACACTGTTGTAT
GGTTCTGTGGAACGTCCATGTGGTCAGTTTAGAATTTCAAGACTGATAACAATTTTGGTATTTTTTAGGG
GTAGGTATGTCGTAGATTTTACATGGAGAACGCGGCAAAAAGCGACGTTCAAGTTCTGTGTCA

>Vanabin4 promoter -1793 to +162 atg start codon
TAATACAATAGGAAAGAAACGATGGTATGTCGCTACTGTTGTTGCAATTCTTTATATTACGATCAGTTTA
GTCAACTTGCCGAAGAACTGTCTTTTAAAGCTCCGTTTTCGTATGAATGAATTTAATTTATTACGCCCGA
CGAGGCGGGACAGCGACAAGCGTTGTTGCACAGGGTTTGATTTCATATACCTTGAACCCAATTGCGAACT
TTATAACTTGTTTGTACATAGGTAAACGATGCCGCTGTGGCCGTTTAAAAAGTGTACACTTTTAGGACGC
GCTGTATATTATATTTAGCGGAGGACTTGCGCCCCCTGGCGGCCAGTTTTTTTTAATTTAGAAAAAAACG
AGCGGTTTAAAATTAGGTTTAAGTTAAAAATACAGAGCCAGATACCGCAGTTGGAAAAAAACTGAAAGCC
AATACATTGTAGTTTTACATTGTTTTTTTTTGTTTTTTTGCAGGATCATAAAATTTCGTTTGGAGCTGCG
AAAAAATATTTCAATGCACAAATTGTTCCGATGCTTAAAAGTACGATGAAAAACACCGATCTATATCATT
AGACATCAAAATCATGAGAATAAATGTAACTTGCTTTATCCTGGCGTGGCCGGAACACGACAGCCGTTGT
AACACGGACGTATAGGCTACACTCAATGTCAGCTTATGAGTAACCATGTATGTTACTTAGTGAGAGTATA
TTATATACACTATCGTCTTCGCCAGTTTAAGACTACTGCTATTAACTTTCTTCAATATATTTTCCGCAGC
AGGTTTTGGGATTCTTTCGTAGATGTGCGTTAACGTTTAATGTACGCTTGTGGAACTTGTTTGCCAGTAA
CAATGCAATTTAACCAATGTCAAAACAAGCGCCTGCATTTTGCAAAACAAAACCCAGCGGCGTCTGAGCG
GTTCGAGCTCGAGCACCGCAGACTAAGATTCTGTGGGTATGTTTTCGACTCAATAACCCCCACCAGCTCA

AAACAACAATATTTTCCTCCTTTAATTCTCTTCAACGTTTCTTTCTTTTATTCCACCACCGACGTGAAGA
GAGAAAAAACGCAGGAGGTGGTTCGAACATTCTAGCAAACGCGGGTCAAACGAGGCAGCAAGACCAACGT
TTTTTTTTGTGGAAATATACTCGCCTAAAACAGCATCTTCCTTCACGGCAAAATATGATTCTTTTTAAAT
AACAGAGATGCAATTTTGGATGAAAAATATATTCAACAATATAAAAAACGTTTGGAATTTTGTCGGTTGG
ACATTTCGTAGCATTTGATCCTGAATGTTATTTCGTTACGAGAATCGTGCAACTTATAAAGGAGACCAAG
GACAGATTAATGAAATATATTACAGTTAGCATGTGAAACAAAGCGAACTCTCCCATATACCGTGTTGAAT
AAGGCGACAAACAGAAAAGTTCAATTGAGGCCAATTTATGAACGAAAACGTCAAAAACAAGTGCATTGTA
CCCCGTTGACTGATGTTTTACCAAAAGACAAAAAACGAACCATTTTCGCTTCACTGAAGCTCGACGCTTT
GAACGCGTGATCTTTTGACAAATTCCATCATCAGCAGAAACAGCTAGCGGAATTGGAGCACAAGGTTAAT
CCAACCGTGCAGAGCGATTTACTTAGCTTGCGTTTTGCAAGGAAAAGCTTCATTTGGAGTCAACAGGACT
GTAAGTAGTTGCTGTTGTTTGTAGCTAATGCTAATTGTTAAACAGGGGAGCGAATGTATTTAAAATAATA
ATTATTTTAATTATCTACATGCAGAAGCTATCAACCGATCAAAATGAAAACGTTCTGTGTTGTTACAATT
GTACTTGTGCTTGCATCGGTGTGTGTTGATGCCCGTGGAAACCGCCATCATGGAGGATTGATGGGAACTG
GGGTGCCAAGGTGCCTGAAAACGTGCAAAGATGATTGCACGGAAATGAAACCTTGCGCTTTAGCT

>Eef1a promoter -1955 to -1
GTGACGGGAAAACGATAGTCGTTATAACACGAGTATTCGTACACCTCGTGCGAGCTAACGAGCTACCATA
TATGTTGTGGGCGAATAAAGGTTTTATAAATATAACATTGGTTTTATAAATAAAACAACGCCATTTTAAA
GTCGGTTACATAATTCTGTAACTAGTTCAAATTGAACGGTAAACGTAAATAAAAACCTTGACCGTCTTAC
CCAATTATATAAAAACACTTTGAACGCTTTTTAAGATGGAAGGGTATGGCCATGCCTAGATAATTCTGTG
GACCATCTCACCCCAACCTATTACAGAACGGTCGTAATAATGAAAATGGGTACCATTTTTAGGCATATAG
ACTGATTCCTCCTTTCTAGAAACGTAAGCAGTATACACAGAAAAAATGAAGTGTGATTCTGTGCAATTAA
ACCGTTCTAAATTCATAGCCGACTGAATTTCTAATTAAGTGAATGTCTGACCTAGATTTATTGTTAAGTT
TAGCACCAAATCTGAGCCAGCGATAAGCAGTCTAATTAAATTGGCTGCTGGCGATAAAATAGGTCATCCT
GAAAAATCGTTTGCGCCTTTATTTAAAATATAGTAGAGTGGGGAAAGACGGGACATCTTATCGTTCTATT
TTCTCGTCCCATTTCGTAGTAAACAAAGAACATTCAAAAAATATAAAACCATAACTTCAAAACTTCAATA
GACCGTTGTCAACTGTTTAAAACACAATAAGAGAATTTGGATATTATGTGCTAAAGGTGTCCCATCTCCC
CCCACCCTACTATATCTGTTTATAGTTCTGTGGGGTAAGATGAGATACCGTTAACACCTAAACATTTTTA
CTTTAAACAATCAACCACGTTTTTTATAGTCGTAATGGACATGTGGTTACATAATTCTGAAAATATTTTT
TGCCCCCGACCAAAAGACGCGAAGAGTAAAAACATGTCTCAGCTTATATTCCCCACATAAATATATTTTT
GTACTGTTTGGTGAATTTATAAACTTATATTACCATGCATATACGTTATGTTACTGGTATTTTCTCAGTA
GGCAAATTCATTTGTCCACGTTTTATAGGTTTTCAATATTTATGATTTTTAAAATGCTAAAAATGTGGGA
GGGGGGTTGAAAGTACAATACAAACACACAAAACAACTCAAACTAAAGATTTATAGTTATGCTAATTCAC
CTACACAATATAACAAGATGTGTAATGCAACCATGTGTTTATGATGAGCGCTAACATATTTTGTAACCAC
TCAAATTCCCCGCCACACGAGGATAATGAATAGGTGACTCTGTAGTCTGTACATCTTAGACTGAAATAAA
GATTATAAATCTACGAAATAAAATAATTTCTGCTCACTGATTATACTTCTGTTTTATAGATTAGAAACCG
TTTCTAATAAATGACCTAATTCGCTATACACACACGCTGTGCGCGAGATAATCATTCTCGCACCCCGTTT
ATTGTGTTAAAATTGCCGCCTAGATTCACAAAGCGTGACGGCTAGAGCCAGCAACGTGTCGCCTTCAATT
ACGCAACATCCGGGTTGCGCAATTCTGGATATAAAAGAACTAACAAAGATGACGTAGCTACCTTTTTCAG
TTCAGACTTACGAAAGACTCACGTGTCGGCGGTCTACTTGTCCTTTTCGAGCTGTGGCAATTTGGTGAGT
GGTTCTATCTTATATCTGAGTACATCTCTAAGGAATTATAGTTTGATTAGTTAAGTTTTTATTGTTAGGA
AAGATGAAATCATTAGGTTTTACTTAGTTTAAGTATGTTAGTACTGGTTAGGCGTTTGAATTATTGAAAA
ACTCAGTTCGTTAACTGTAGTAGTTCTGGTAGCTTAGCAAGTATACCCTGTATACGCCTTTTGGCTTTTT
AACAATAACTTAAACTTATTTTACAGCAAATTTCTGTGCATTCGGTTAACCCCAACCTTCCAAA

>Cas9::CionaGeminin-Nterminus (from Song et al. 2022[8], which originally described it as having the Geminin sequence from human instead)
nls::Cas9::nls (described in Stolfi et al. 2014)
Ciona robusta Geminin N-terminus

ATGGCTAGCCCCAAAAAGAAGAGGAAAGTGGACAAGAAGTATTCTATCGGACTGGACATCGGGACTAATA
GCGTCGGGTGGGCCGTGATCACTGACGAGTACAAGGTGCCCTCTAAGAAGTTCAAGGTGCTCGGGAACAC
CGACCGGCATTCCATCAAGAAAAATCTGATCGGAGCTCTCCTCTTTGATTCAGGGGAGACCGCTGAAGCA
ACCCGCCTCAAGCGGACTGCTAGACGGCGGTACACCAGGAGGAAGAACCGGATTTGTTACCTTCAAGAGA
TATTCTCCAACGAAATGGCAAAGGTCGACGACAGCTTCTTCCATAGGCTGGAAGAATCATTCCTCGTGGA
AGAGGATAAGAAGCATGAACGGCATCCCATCTTCGGTAATATCGTCGACGAGGTGGCCTATCACGAGAAA
TACCCAACCATCTACCATCTTCGCAAAAAGCTGGTGGACTCAACCGACAAGGCAGACCTCCGGCTTATCT
ACCTGGCCCTGGCCCACATGATCAAGTTCAGAGGCCACTTCCTGATCGAGGGCGACCTCAATCCTGACAA
TAGCGATGTGGATAAACTGTTCATCCAGCTGGTGCAGACTTACAACCAGCTCTTTGAAGAGAACCCCATC
AATGCAAGCGGAGTCGATGCCAAGGCCATTCTGTCAGCCCGGCTGTCAAAGAGCCGCAGACTTGAGAATC
TTATCGCTCAGCTGCCGGGTGAAAAGAAAAATGGACTGTTCGGGAACCTGATTGCTCTTTCACTTGGGCT
GACTCCCAATTTCAAGTCTAATTTCGACCTGGCAGAGGATGCCAAGCTGCAACTGTCCAAGGACACCTAT
GATGACGATCTCGACAACCTCCTGGCCCAGATCGGTGACCAATACGCCGACCTTTTCCTTGCTGCTAAGA
ATCTTTCTGACGCCATCCTGCTGTCTGACATTCTCCGCGTGAACACTGAAATCACCAAGGCCCCTCTTTC
AGCTTCAATGATTAAGCGGTATGATGAGCACCACCAGGACCTGACCCTGCTTAAGGCACTCGTCCGGCAG
CAGCTTCCGGAGAAGTACAAGGAAATCTTCTTTGACCAGTCAAAGAATGGATACGCCGGCTACATCGACG
GAGGTGCCTCCCAAGAGGAATTTTATAAGTTTATCAAACCTATCCTTGAGAAGATGGACGGCACCGAAGA
GCTCCTCGTGAAACTGAATCGGGAGGATCTGCTGCGGAAGCAGCGCACTTTCGACAATGGGAGCATTCCC
CACCAGATCCATCTTGGGGAGCTTCACGCCATCCTTCGGCGCCAAGAGGACTTCTACCCCTTTCTTAAGG
ACAACAGGGAGAAGATTGAGAAAATTCTCACTTTCCGCATCCCCTACTACGTGGGACCCCTCGCCAGAGG
AAATAGCCGGTTTGCTTGGATGACCAGAAAGTCAGAAGAAACTATCACTCCCTGGAACTTCGAAGAGGTG
GTGGACAAGGGAGCCAGCGCTCAGTCATTCATCGAACGGATGACTAACTTCGATAAGAACCTCCCCAATG
AGAAGGTCCTGCCGAAACATTCCCTGCTCTACGAGTACTTTACCGTGTACAACGAGCTGACCAAGGTGAA
ATATGTCACCGAAGGGATGAGGAAGCCCGCATTCCTGTCAGGCGAACAAAAGAAGGCAATTGTGGACCTT
CTGTTCAAGACCAATAGAAAGGTGACCGTGAAGCAGCTGAAGGAGGACTATTTCAAGAAAATTGAATGCT
TCGACTCTGTGGAGATTAGCGGGGTCGAAGATCGGTTCAACGCAAGCCTGGGTACCTACCATGATCTGCT
TAAGATCATCAAGGACAAGGATTTTCTGGACAATGAGGAGAACGAGGACATCCTTGAGGACATTGTCCTG
ACTCTCACTCTGTTCGAGGACCGGGAAATGATCGAGGAGAGGCTTAAGACCTACGCCCATCTGTTCGACG
ATAAAGTGATGAAGCAACTTAAACGGAGAAGATATACCGGATGGGGACGCCTTAGCCGCAAACTCATCAA
CGGAATCCGGGACAAACAGAGCGGAAAGACCATTCTTGATTTCCTTAAGAGCGACGGATTCGCTAATCGC
AACTTCATGCAACTTATCCATGATGATTCCCTGACCTTTAAGGAGGACATCCAGAAGGCCCAAGTGTCTG
GACAAGGTGACTCACTGCACGAGCATATCGCAAATCTGGCTGGTTCACCCGCTATTAAGAAGGGTATTCT
CCAGACCGTGAAAGTCGTGGACGAGCTGGTCAAGGTGATGGGTCGCCATAAACCAGAGAACATTGTCATC
GAGATGGCCAGGGAAAACCAGACTACCCAGAAGGGACAGAAGAACAGCAGGGAGCGGATGAAAAGAATTG
AGGAAGGGATTAAGGAGCTCGGGTCACAGATCCTTAAAGAGCACCCGGTGGAAAACACCCAGCTTCAGAA
TGAGAAGCTCTATCTGTACTACCTTCAAAATGGACGCGATATGTATGTGGACCAAGAGCTTGATATCAAC
AGGCTCTCAGACTACGACGTGGACCACATCGTCCCTCAGAGCTTCCTCAAAGACGACTCAATTGACAATA
AGGTGCTGACTCGCTCAGACAAGAACCGGGGAAAGTCAGATAACGTGCCCTCAGAGGAAGTCGTGAAAAA
GATGAAGAACTATTGGCGCCAGCTTCTGAACGCAAAGCTGATCACTCAGCGGAAGTTCGACAATCTCACT
AAGGCTGAGAGGGGCGGACTGAGCGAACTGGACAAAGCAGGATTCATTAAACGGCAACTTGTGGAGACTC
GGCAGATTACTAAACATGTCGCCCAAATCCTTGACTCACGCATGAATACCAAGTACGACGAAAACGACAA
ACTTATCCGCGAGGTGAAGGTGATTACCCTGAAGTCCAAGCTGGTCAGCGATTTCAGAAAGGACTTTCAA
TTCTACAAAGTGCGGGAGATCAATAACTATCATCATGCTCATGACGCATATCTGAATGCCGTGGTGGGAA
CCGCCCTGATCAAGAAGTACCCAAAGCTGGAAAGCGAGTTCGTGTACGGAGACTACAAGGTCTACGACGT
GCGCAAGATGATTGCCAAATCTGAGCAGGAGATCGGAAAGGCCACCGCAAAGTACTTCTTCTACAGCAAC
ATCATGAATTTCTTCAAGACCGAAATCACCCTTGCAAACGGTGAGATCCGGAAGAGGCCGCTCATCGAGA
CTAATGGGGAGACTGGCGAAATCGTGTGGGACAAGGGCAGAGATTTCGCTACCGTGCGCAAAGTGCTTTC
TATGCCTCAAGTGAACATCGTGAAGAAAACCGAGGTGCAAACCGGAGGCTTTTCTAAGGAATCAATCCTC
CCCAAGCGCAACTCCGACAAGCTCATTGCAAGGAAGAAGGATTGGGACCCTAAGAAGTACGGCGGATTCG
ATTCACCAACTGTGGCTTATTCTGTCCTGGTCGTGGCTAAGGTGGAAAAAGGAAAGTCTAAGAAGCTCAA
GAGCGTGAAGGAACTGCTGGGTATCACCATTATGGAGCGCAGCTCCTTCGAGAAGAACCCAATTGACTTT

CTCGAAGCCAAAGGTTACAAGGAAGTCAAGAAGGACCTTATCATCAAGCTCCCAAAGTATAGCCTGTTCG
AACTGGAGAATGGGCGGAAGCGGATGCTCGCCTCCGCTGGCGAACTTCAGAAGGGTAATGAGCTGGCTCT
CCCCTCCAAGTACGTGAATTTCCTCTACCTTGCAAGCCATTACGAGAAGCTGAAGGGGAGCCCCGAGGAC
AACGAGCAAAAGCAACTGTTTGTGGAGCAGCATAAGCATTATCTGGACGAGATCATTGAGCAGATTTCCG
AGTTTTCTAAACGCGTCATTCTCGCTGATGCCAACCTCGATAAAGTCCTTAGCGCATACAATAAGCACAG
AGACAAACCAATTCGGGAGCAGGCTGAGAATATCATCCACCTGTTCACCCTCACCAATCTTGGTGCCCCT
GCCGCATTCAAGTACTTCGACACCACCATCGACCGGAAACGCTATACCTCCACCAAAGAAGTGCTGGACG
CCACCCTCATCCACCAGAGCATCACCGGACTTTACGAAACTCGGATTGACCTCTCACAGCTCGGAGGGGA
TGAGGGAGCTCCCAAGAAAAAGCGCAAGGTAATGGCCACGAAAAATATTCTTCAAAATATAAATGCACAA
TGGAAGGAGAATGACAACAGATCACCAAGTAGAAAGCGACGGTTAGATGACGTCACTGAAGAATCACAAT
TACCTTCCACGACCAAACGACGTCATCTTCAAACAAATACAAACGTTGTAAATTCCACAGGATTGAAACA
AGGCCTGACAAATGTGAAAAATTCAATAAATCCAAAGAACAAATCAATAAAAAATTTCTTTTCTGATATT
CCACGTGTGTCATGTACTAAATCTGAAAAGATTCAAATTTTTAAAGAAGCTAAGAAAACTCCAAAAAAGA
ATGCAACCACTCAGACAAGGAGTGAAGCTGAAGAATTGGTCTGCAGTGATCAACCCAGTGAAAAATATTG
GGAACTCTTAGCCGAGGAGCGAAGGAAAGGGTTGTAA

>Pax2/5/8.a tv1 “rescue” coding sequence, sgRNA sites mutated
ATGAACTGGGGATCAGCAATGGCGGTTGGACCATCCAGTGTGGGACATCCATTCGTGGGATCAGGAATGG
CGGCCTCACTCACCCCTTCAAGATCAGGGCACGGTGGGGTGAACCAGTTGGGCGGGGTTTACGTGAACGG
CCGACCATTACCCGACCAAGTCCGACAACAAATtGTcGAtCAgGCtCAtATAGGGGTTCGACCTTGcGAt
ATaGCacgcCAgctCCGGGTGTCGCATGGTTGTGTAAGCAAGATATTAGCAAGATATTACGAGACAGGCA
GCATCCGACCTGGTGTaATtGGaGGttccAAgCCaAAaGTtGCaACtCCaagaGTtGTGGAGAAGATATG
TGATTATAAACGACAGAATCCAACTATGTTTGCTTGGGAGATACGAGACCGATTGTTGAGTGAGGGAATT
TGTGACCATGATAATGTACCCAGTGTTAGTTCGATCAATAGGATTGTCCGAAACAAAGCTGCAGAAAATG
CAAAGTCGCACCAACAACTCATGGTCCCAATGACGCCGTCATCGCTGGACAACTACAGCGAGCACATCGG
GCAGGTGTCGCGCATGAACGGTTTCAATAGATTCGCTTCATCGACCACAGCATTACCCACAATGCAACAA
TCAAGTCATGTGACTTCTCAGATGAATTTCGTTCGTAAAGAAAACAAGGGATTGGATTATTCGTACGATT
GTCGCGGTTCAACGAAcTCACCCAACGTAGCTACTTACCCAGTGTTACCTCACAATCAACCTCGAGCTAA
CTGTGATGTCACAATCAGCCCAATGACATCACAAACCAACACTGCAAACGCGACAGTTTCACCCAGCAAC
AGTGGGGGTTACTCTGGGTCAAGTTTTGCCCCGATCACAACCGCGTACGCTCCGACAGGTGAGTTTGTGG
ATTATGGATATCAACAATACAATCAACACTGGAAGTTTGGCCAACAGCACCATAATGATAGCAACACTGG
TAAAGTTTTGAATCTTCGAGGTAAAGAACATCCGGCAACACTGGAGATGGTCAGCGCTCAATAA

>Pax2/5/8.a tv2 “rescue” coding sequence, sgRNA sites mutated
ATGAACTGGGGATCAGCAATGGCGGTTGGACCATCCAGTGTGGGACATCCATTCGTGGGATCAGGAATGG
CGGCCTCACTCACCCCTTCAAGATCAGGGCACGGTGGGGTGAACCAGTTGGGCGGGGTTTACGTGAACGG
CCGACCATTACCCGACCAAGTCCGACAACAAATtGTcGAtCAgGCtCAtATAGGGGTTCGACCTTGcGAt
ATaGCacgcCAgttCCGGGTGTCGCATGGTTGTGTAAGCAAGATATTAGCAAGATATTACGAGACAGGCA
GCATCCGACCTGGTGTaATtGGaGGttccAAgCCaAAaGTtGCaACtCCaagaGTtGTGGAGAAGATATG
TGATTATAAACGACAGAATCCAACTATGTTTGCTTGGGAGATACGAGACCGATTGTTGAGTGAGGGAATT
TGTGACCATGATAATGTACCCAGTGTTAGTTCGATCAATAGGATTGTCCGAAACAAAGCTGCAGAAAATG
CAAAGTCGCACCAACAACTCATGGTCCCAATGACGCCGTCATCGCTGGGTATCCACACAAACGGTCCGAT
ATTAATGGAACACGAGATTCGCGGAACGTACACGATTAACGATATATTAAGGTTACCCCACAACCCCCTT
CCCCCACATAACCCCCCCACCACTGAAACAGACCCAACACATTATAAACAAGCCGAAAATGGGATCCATT
ATAATCATAACGACAACTACAGCGAGCACATCGGGCAGGTGTCGCGCATGAACGGTTTCAATAGATTCGC
TTCATCGACCACAGCATTACCCACAATGCAACAATCAAGTCATGTGACTTCTCAGATGAATTTCGTTCGT
AAAGAAAACAAGGGATTGGATTATTCGTACGATTGTCGCGGTTCAACGAAcTCACCCAACGTAGCTACTT
ACCCAGTGTTACCTCACAATCAACCTCGAGCTAACTGTGATGTCACAATCAGCCCAATGACATCACAAAC
CAACACTGCAAACGCGACAGTTTCACCCAGCAACAGTGGGGGTTACTCTGGGTCAAGTTTTGCCCCGATC

ACAACCGCGTACGCTCCGACAGGTGAGTTTGTGGATTATGGATATCAACAATACAATCAACACTGGAAGT
TTGGCCAACAGCACCATAATGATAGCAACACTGGTAAAGTTTTGAATCTTCGAGGTAAAGAACATCCGGC
AACACTGGAGATGGTCAGCGCTCAATAA

>caBMPR1b coding sequence, gac (Q202D) constitutively active mutation
ATGGCGGCCGCAACCATGCTGGACAATGGACTACCTCGTATGCAGTGTCGTTGTATTGGAGAATGCCCGG
ACCACAAATACAACTCCACATGCGACCCGAAACCCAACGCAAAGTGCTTCAAAAAGTTGTACATTAATGA
ATATGGTGAAGAGGAGTTACGAGCTGGGTGTTTAGGTTCACAAGATGATTACTTGAACCAATGTCATAAT
AAAGCAAAAACAGAATTCGAAGTTCCAACTGCAGTGGCTTGCTGCAACAATGGTACAATGTGCAACGATT
ATTTAGACCTCGGGTTGCCTGAATATTATGATGAACCATCTGACACACCAGTTTCAGAAGCAAATTCGGA
TGTTATCACCGTTATAGCTGTTACAGTGCCAGTCTTCTGCTTTTTGTTTGGTTTAATAATAATGTTTTAT
TACATAAGATTATGCCGTAGAGAATCGCTACGACGACGTCAAATTGAGAATGAAAACAAAAAAGCTCTTT
GTCCTGTTTTATATGGAGATCGGGGAGACTTGGAAGGACATAATGAGATGATGGAAAATTGGTCCTCACT
TGCTGGTACATCTTCAGGATCAGGGATGCCTCTACTTGTACAGAGAACGATATCACGAGACATCGAAATT
CTTCACGAAATTGGAAAAGGAAGATATGGGACAGTGATGCTGGGGAAATGGAGAGAAGAGAAAGTTGCTC
TTAAAATATTTAATTCCAGTGATGAAGAAAGCTGGTTTCGAGAAACTGAAATTTACCAAACTGTGTTGCT
ACGTCATGACAATATATTAGGTTTCATAGCAGCCGATATATCTGGTGCTGGCTCATGGACTCAGCTGTTC
CTAATTACAGAGTACCACAAGCATGGCTCCTTATACTACTACCTACAAAACAGAGCCATCAACATAGCAG
AAGCTCTTAAACTTGCTTATACAGCTTGTTGTGGGTTAGCACATCTACATACAGAGATAGCTGGCACACA
AGGCAAGCCAGCCATTGCACACAGGGATGTAAAATCGCAGAATATTCTCGTAAAACTTGATGGACAATGT
TGTATTGCAGATATGGGACTGGCTGTCTGTTTCTCCAGATTACATGAAACGATTGATGTTGGAAAGCATG
ACCGCTCAAGAAGACAAGGTACCAAGCGTTACATGTCACCTGAGGTGCTTGCCCAGTCTTTTCACCCAGA
CTCCTTTGAAGCATACAAAGCATCAGATGTGTACAGCTTTGCTTTGGTGTTATGGGAAATAATCAACAGG
ACTGAAGTCAACGGATTTGCAAATGACTACCATCTTCCCTACCACGATGTGGTGGGTAATGACCCTGATT
TTGATGAGATGAGAAAAATAGTTGTTCTTGAAAATCTCAGACCAGAGATTTACAAACAGTGGCAAGCACA
TAAGATAATGTCAACATACACTACAACCCTGCAAGAATGTTGGTCCCCTCGACCCGAGTCTCGTCTCTCA
ATGCTCCGTCTCCGCAAGACTCTCTACTTTCTCCTCAATGGTGGACGAGGCCCAGTGGAGGCGGGAAGCT
CTGAAAGTGACCGAAAACCGTCTGCCTCTTCCAGCTCTTCTGTTAAAGAAGAGGAGCATTCAAGTTGCTA
A

>dnBMPR1b coding sequence, early STOP dominant-negative truncation
ATGCTGGACAATGGACTACCTCGTATGCAGTGTCGATGTATTGGAGAATGCCCGGACCACAAATACAACT
CTACATGCGACCTGAAACCCAACGCAAAGTGCTTCAAAAAGTTGTACATTAATGAATATGGTGAAGAGGA
GTTACGAGCTGGGTGTTTAGGTTCACAAGATGATTACTTGAACCAATGTCATAATAAAGCAAAAACAGAA
TTCGAAGATCCAACTGCAGTGGCTTGCTGCAACAATGGTACAATGTGCAACGATTATTTAGACCTCGGGT
TGCCTGAATATTATGATGAACCATCTGACACACCAGTTTCAGAAGCAAATTCGGATGTTATCACCGTTAT
AGCTGTTACAGTGCCAGTCTTCTGCTTTTTGTTTGGTTTAATAATAATGTTTTATTACATAAGATTATGC
CGTAGAGAATCGCTACGACGACGTCAAATTGAGAATGAAAACAAAAAAGCTCTTTGTCCTGTTTTATATG
GAGATCGGGGAGACTTGGAAGGACATAATGAGATGATGGAATACCCATACGACGTGCCAGACTACGCTTT
ATAA

>dnFGFR (dominant negative FGF receptor, Davidson et al. 2006[9])
ATGATACAACTACAAAATACGTTTATTTTTATCGCTTTGACAATTTTTACTTCTGCTTCAACAACAAGCT
TAAAGAATGAAACCAAACCCCTCAACACAATTTCAACGCTAGCTGCTCAAACAAACATTTCAAACCCAGA
AGACGATTTGTTCGATACAAACGGAGCACCAAAAAGTGATACTGTGAATGCATCTACAACTACGGATCGT
CACAAGATTCCACGCTGGGTCAATGAACAGAAGATGCAAAAGCGACTTCACGCTGAACCAGCAGGTAACA
CTGTCCAGTTTAGATGTGCAGTTCAAGGTGCAAGACCAATCACCGTGGATTGGTATAAAGATGGGGAACC
AATCAAGAAGAATGGAAGACTGGGGGGGTACAAGTTCCGTCAACGCAACCAGCAAATATCATTGGAGTCG
GTGATAATGTCCGACCGTGCTAAGTACATGTGTGTGGCCCATAATAAGTACGGCTCCATCAACCATACTT

ACGAACTGGATGTAGTAGAACGTCTGGCACACCGCCCCATCCTCCAATTTGGATTACCAGCAAATAAAAC
TGTCAAGGTGGGAGAGGATGTGACCTTCAAATGCAAGGTCTACAGTGACCCTCACCCTCATATGGAGTGG
TTAAAGCATGTAGAGGTTAACGGGTCAAAATACGACCCAGTTTCTAAGTCCCCCTATGTGATTACATTGA
AGAGAGCTGGTATTAACACAACTGACGCGGAGATGGAAAAGTTAACATTGAAAAATGTTTCATTTGCCGA
CGCTGGGGAATATACTTGTCTTGCTGGAAATTCTATCGGGGTGTCTCATGTTTCTGCATGGCTCACAGTG
CTGCCAGTGGTAGATGAGAATGATGTATGGACCGAAGAAATCCCACAAGACACTCATTATCTCATATATA
TCTTTGGGGTTGTGTGCTTCATAATACTACTTGCGTTCATTGTGTATATGTGCAACTCTCGCTATCAAAA
TAAAGATCCTCCCCGTTTGATCCCGATCGAGAACCCCGACAACATCCCCCCCATGTCGAAGATGGAGGAG
CCGGTGATGTTGTTCGGGAACTAA

>dnEph.c (dominant negative Eph.c receptor, Picco et al. 2007[10])
ATGTTTTTTCTATTTATCGTCTTCTACTGCATTTCAATTGTAACTGGAGAAATACACGTTTTATACAACA
CTAAAGTTGCGACAAGTGATTTAGATTGGGCCTTACACCCCACATATGGAGCGTGGGAGGAATTGAGCGG
ACTCGATGTAGACGGTAACACAATACGTTACCACCAAGTATGTAACACAGGAATGGATGAACAGGACAAT
TGGGTACGATCCCCATTTATCGACGCAAAGTCGGCCCAGCGTATTTATATGGACATCGAATTTTCGGTTA
TGAAATGTGAGGAATGTCGCGAAACATTCGCGTTGTATTATTACCCGTCCAGTTCCGACACTGCTACCAC
TACATTCCCGCCTTGGAGAGAAAACCCCTATATCAAAATCGATACATTAGCAGCTGGTGAAAGATTCGAT
TCCGAAACCGTAGGTGCGGAGGGAATTAATAAGAAAACATTGGTTATTGGACCATTATCAAGACGCGGAT
TTTATATCGCTGCTCAGGACCAGGGCGCCTGTATGTCAATTATGAGTTTGAAATTGTATTATTACCACTG
TGAGGAAACGACCCATAACCTGGCTTATTTTCCGAACACCATATCGGGCGGCGGTATCGCTGAATTGGTG
TCGCAATCTGGAAAGTGCGTTTCGAACTCAGTATACGCGAATGAAGTTCCGAAATACCGATGTAACATTT
ACGGCGAATGGCAAGTCCCTACTGGTTCTTGTCAGTGCAGAGCTGGTTATGAACCAAATACACAACTCAC
CGCATGTACAGGCTGCATGGTTGGCAAGTACAAATCTACAAACGGCAATACACCTTGTCAAGTCTGCCCA
CAGCACAGCGTTACACATTCAACATCCGCTAGTCACTGCACTTGTGTGGCTGGACATTATCGAGCCGAAA
ATGACCCAATCAGCCAAGCCTGCACTCGGCCCCCCACTAAGCCTCGGAATGTGACTCATGTTCAAAACAA
GACTTCTCTCTTATTGTCATGGGTCCCACCTTCAACTACAGGGGGTAGAACTGATATATATTACAGCATC
TCATGTGAACTGTGCGATTCTGAGCATGAGAACTGTCAGCCCTGCAATGTTGATGTTCAATACAGACCAA
GCAACCACCAGCACACCACAACAACGTACATGGAAGTCTCAAATCTTAATCCCTTCTCATGCTATAAGTT
CAAGGTTTCTTCTTCAAATGGTGTTTCTAGGGTCAGTGTGGAGCCTGAACAATTTGAGCTAATAAAGATT
TGCACCAACGCCGCTGCTCCGTCTGCTGTCACGGGCCTTAAGTTTATTTGGATTGGGGAAGTTTCGGCAA
CTTTAAGTTGGTTGCCGCCCACACATAGCAACAAGATTGTTGGATATGAAGTGCAATTGTTCCAAAACAA
TCAGCGTTTAAAACTGACAAAAATTATGGAGGTTTCAACACCAAACGTTACAATAAATGGTTTGAACCCT
GGATGGAAATATGTAGTGATGGTACGAGCATGTAATAACGATGGGTGCGGCCAGTTCAGTGAAAAACTGC
AGCTAGTTACTTATGACAAAGGTTCTGAGCCTGAATCTATAGTGGAAGGCAGTACTACATGGATTGGAGG
AGTTATTGGTGGGGTTCTAGTTATCGTTATAATAATTATCGTCGTGATGATAAAAAGACGACAAACCGAT
AAAAAGAAACGAAAAGAAATCCAAGCCAGAACAAAACTCAACGAAAACACTCAACAACTAAACCAGAGTA
GCTTTTTACAAACCGCAGGGCGAACTTATGTTGATTACCGTGACCCCCACAATGGGGTAAAAGAAATCGC
AACCGAGATTGATCAAACGAGAATTAAGATCGACAGTGTTATTGGAAGAGGTGAGTTCGGTGAAGTGTGT
CGTGGTAAAATGTTGACGGGCAAAACAACGACTTCTGTCGCCGTGAAACGATTAAAACACGGAGCAAGTT
TAATTGACCACACCAACTTTCTACGAGAGGCATGCACTATGGCACAGTTTAAAGACCCGAATATTATTCA
ACTGAAAGGAGTGGTTACTAAAAGCATCCCCGCAATGATAATCACGGAATTTATGGAGCATGGTTCGCTT
GATAAATTTTTACAGGCTCGTTCTGGGCAGCTTACTGTTCTACAATTACTTGAAATGTTACGCGGCATCG
CAAGTGGAATGAAATATCTTTCATCAATGAAATATGTTCATCGAGATTTAGCTGCAAGAAATATCCTCGT
TAATTCTCAACTTGTTTGTAAAGTATCCGACTTTGGTCTTTCAAGAACTCTTGAGAATGACCCTCAAGCT
ACCTATACCACACAGGGTGGAAAGATTGCTCTCCGCTGGACAGCCCCTGAGAGCATCCGTTGTCGTCAAT
TTACATCAGCAAGTGATGTATGGAGCTATGGGATCGTTATGTGGGAAGTCATGTCTTATGGGGAGAAACC
TTACTGGGATATGAGTAATGAAGTTGTGACAGAGGTATTAGAAGATGGATACAGGCTCCCGTCACCTGAG
GGTTGTCCTACTCCTGTCCATAGTTTGATGTTGAAATGTTGGTCTTATGAACCGAAACGCAGACCAACCC

TGCTGGAAATTATTAAGACTTTGGATCATTTTATTAAACAGCCCAGCTCTCTTCAAGACGACATGGAAGC
TGATGCAAGTGCCCCGCTGTTAAAGCCAGACAGTCCCAACAGCATTCAAGATGTATCAACATTAGACGAG
TGGTTAGACATGGTAAAACTTGGAAGATACCGCAGGAGTTTTCATAACAACGGAATTAATGATCTTGAAA
GCTTGGCTCACATAAGTGAAAGTGAGTTGGACAGACTTGGCATAGCGTCACCTTCCCACCGCACCAGACT
ACAGGGAGGCATTAACACTTTACGACAACATTTGGTCGAAGTAAACGAAGTTCAATCCAGCAGTAACGCA
CCTTACGACGCCCACGCTTCAACGCTACCAATAGCTCACGGGAAACCAAACAATCCTGTTGCAGTTTGA

>caMEK (from Razy-Krajka et al. 2018[11])
ATGCCTCCTAAACGTAAGTTAAACCCGTTGAACCTAACACTTGAGGGGAGTTCGCCTATCCCCAATCAGC
AGGTCTCTCAGTTCATTAAGCAGAAGCAAAAAGTGGGAGTCATGGAAAATGCTCAGAACTCTGACTTCAC
AAAGAAAGGAGAACTGGGTGCCGGAAATGGGGGTGTTGTTCATCTTGTGGTGCACAACGCAACTGGCTTT
GTCATGGCGCGGAAACTTATTCATTTGGAAGTGAAGCAAGCCATCTTGAACCAAATCACACGAGAATTGC
AGGTTTTGCATGATTGTAGAAGCCCGTACATTGTTGGGTATTATGGACTTTTTACAGTGATGGGGAAAT
CAGTATTTGTATGGAAAGCATGGACGCTGGGTCGCTTGATTTAGTGCTGAAGAAGGCGCGAAAAATTCCT
GAGATTTACTTGGGAAAAGTGAGCAAAGCTGTTATCCTTGGCCTCAAGTACTTAAGAGAAGAACGTAGCA
TCATTCATCGCGATGTAAAACCATCAAATATTCTCGTTAATTCCCGAGGAGAGATTAAGTTATGTGATTT
TGGCGTGAGCGGGCAACTGATCGACGATATGGCCAACGAGTTTGTAGGGACAAGATCATACATGGCCCCA
GAACGCTTGCAAGGATCCAAGTATACAATCCTTTCAGATATCTGGTCTCTTGGTCTCTCACTTATTGAAA
TGGCAATTGGTAGATTCCCTATCCCGCCACCCACAGCCAGCCAAATAGCAGCCATATTCAACACTGAAGT
GGCAGGGGGAAGTGGTAAAGCACCAAACCCACATGATGTTGCACGACCAATGGCAATCTTTGAGCTGCTT
GATTACATTGTGAATGAGCCAGCACCGAAACTCCCACAAGGAATTTTCGAGAAAGATTTTTGTGATTTCG
TGGCTAGTTGCTTGAAGAAAGAACCGAAAGAGCGATCAGATCTCGGGGAACTAATGAAGGCTCCATTTAT
TAAAAATGTTAGCTTAACCCAGTATGAGTTTGCTAAGTGGGTTTGCAGTACTATGGGTTTGAAAGCCCCG
AGTCCAGACACTGTACCTGATTAA

>Phox2 complete gene model(new exon 1, new exon 4-7)
ATGGACTACCCTGCGTATTTAGGGGTTGGCACAAATTATGACACGACTGCCTGTATGGTAGCTGCGGCAG
CTAATAACGATCCACACCAGCCTTACGTTACTACTCCATATGGGGACTTTAACTCATGCGCCCACGCCCA
GTCCTCTCAGAGCAACTTCGCGTCAGCTTATACACCGACAGTCCCGATCCGGAACTCCAGTTTTGGTGGC
GGAAACTGCCAGCCACCGATGCCTACAGCGGCCGCGTACGGCCTCAACAGCTTAAGGGATCAGTCGCCAT
ATTCTTCAGTACCTTGTAAGTTCTTCACCGAGACGGCCCATCAACACACCGGGGGCTACGGAGGACTCCA
CGAAAGGAGGAAGCAACGTCGCATCCGGACTACGTTCACAAGCTCCCAGTTAAAAGAGCTTGAAAAAGTT
TTCGCCGAAACTCATTACCCGGATATTTATACAAGAGAAGAACTTGCGCTAAAAATTGATCTCACTGAAG
CCAGAGTGCAGGTTTGGTTTCAAAATCGTCGAGCAAAATGGCGAAAAATGGAGCGAGCAAAACAACAACC
TCAACCCATACATTCTCCTGGGAGTTCCCCTTCTTCTCCCAACAATATCTCGTCCATAAACAACTCGGAA
AATATGATAAGTTCCGCAAGTCCTATGGAAGATATAGCTATTAGCTCAGTAAGCAGTGGAGCAATGATGG
AAGAAACAAACCAAGATCATCATTCAAGTCAGGATGACGTCATGGCTCTTAAAAACTCACTGCTAGGTAG
CGCTGCAGAGCTCGCAAGCATAGTCGGTAGTAAAAAACAAATTAGCACTCCTGTTCGAAACGAAAACATA
ACCGAAAATTCCGGTTTGCTATCTGATGACGTCACCACGCATCAATCCCATCATAATACGTCAGAAATGA
CGTCACCGCCCGGTTTTCGCGGTAAACCAATCCAATCTTTGATGTTACCAAACCATGGACAATCAGGATC
AAGTTATAACAGCAATCAACAGTTCATGCAAGGTCACAACCGATCTCAGCACAACACTTCATCATCGGTG
TCTTGGTTCCTATCCGGCGGGGGTAACAACGTCGCCACCCCGCTAAACACGACTCCTTTCGCCGATGTCC
TATCCGTTCTAACGCGTAACACACCCGGGTGCTCCACCGTCGCCGCTGCGGCAGTTTTGATGACGTCATC
ATCCCCCGTGACGTCAACTACTACGTCATACCAAAGCAGCGTAATGAGAATGAATCCACTGCCGGGGCCT
AATGGTGGATTCGACAAATCGTTCACCAACGGAGTCGGGAACAAACAGAGCAGAAATATGTTTCTGGATC
AGAGTCCAATTAACTCTGGCTCTTGCTCTGCGACAATTAACCACAGCCATAAGTTAAACTGA

>Unc76::StayGold (based on sequence from Hirano et al. 2022[12])

ATGGCGGATCTGCGAGTACCGGACATTCCGCTCGCCTCGTGTGATGATGATGATATCGATAGTAATAAGA
ATTTGAGCAACCATTCATCAGACGAGAAACATCACTGCAACAGCAACAGCGACGAGGAACGTCTTCATGA
CGAGTTCTCTGGATCCCTTGAGGACCTTGTCGGCAACTTTGACGAAAAAATTGCGGCATGCCTGAAGGAC
CACGAGGTGACGACAGCGGATATTGCACCTGTGCAGATACGTACTCAAGAGGAAGTTATGAATGAAAGCC
AAACATGGTGGACATTAACCGGAAACTTTGGAAACATTCAACCTCTCGACTTTGGAACCTCTTCGATATG
TAAAAAGATGGCCGCAGCTCTGGACAGTGATTCATTGAAAGACGACGCATCTACACGCCGAAGTATGACA
AATTCCGATGATGAGGATCTTTTACGACAACAAATGGATGTTCATCAAATGATTGGACATCATCATGGAT
CTACGGATACTGGTGGTGAAACACCTCCACAGACTGCTGATCAAGTTATCGAAGAAATTGATGAAATGTT
ACAGGTACCGGTCGCCACCACTAGTGCCAGCACCCCCTTCAAGTTCCAGCTGAAGGGCACCATCAACGGC
AAGAGCTTCACCGTGGAAGGCGAGGGCGAGGGCAATAGCCACGAGGGCAGCCACAAAGGCAAGTACGTGT
GCACCAGCGGCAAACTGCCAATGTCTTGGGCCGCCCTGGGAACTAGCTTCGGCTATGGCATGAAGTACTA
CACCAAGTACCCCAGCGGCCTGAAGAACTGGTTCCACGAGGTGATGCCCGAGGGCTTCACCTACGACAGA
CACATCCAGTACAAGGGCGACGGCAGCATCCACGCCAAGCACCAGCACTTCATGAAGAACGGCACCTACC
ACAACATCGTGGAGTTCACCGGCCAGGACTTCAAGGAGAACAGCCCCGTGCTGACCGGCGACATGAACGT
GAGCCTGCCCAACGAGGTGCAGCACATCCCCAGAGATGACGGCGTGGAGTGCCCAGTGACCCTGCTGTAC
CCTCTGTTGAGCGACAAGAGCAAGTGCGTGGAGGCCCACCAGAACACCATCTGCAAGCCCCTGCACAATC
AGCCAGCCCCCGATGTGCCATACCACTGGATCAGAAAGCAGTACACCCAGAGCAAGGACGACACCGAGGA
GAGAGACCACATCTGCCAGAGCGAGACCCTGGAGGCCCACCTGTAA

**In situ probe templates:**

>Vanabin4 (coding sequence)
ATGAAAACGTTCTGTGTTGTTACAATTGTACTTGTGCTTGCATCGGTGTGTGTTGATGCCCGTGGAAACC
GCCATCATGGAGGATTGATGGGAACTGGGGTGCCAAGGTGCCTGAAAACGTGCAAAGATGATTGCACGGA
AATGAAACCTTGCGCTTTAGCTACATGTCCTTCTGTCTGCCACGCAACTCGCGAAGCGGCCGAAGGCAGT
GGCGCTAATCGCTGCATGATCAGGTGCGGGTTGACTCAATGTTTGCCGAGATTCCCAAGCTGTAAAGCGT
GCGTCGCCCGTTGCGCTGCACCGGTTACCGCATGCAAGCGAAGCAGTTGTGCATCTGAGTGCCCAGCCGG
GATGACCATACCTGAGCACATGCAACTAAGTGGATGCGTTCGTTGCATGAAACGTAATTGCAGAAATATA
ATGAATGGGAACTAG

>Phox2 N-terminus gene model (KH.C14.100)
ATGCCTACAGCGGCaGCGTACGGCCTCAACAGCTTAAGGGATCAGTCGCCATATTCTTCAGTACCTTGTA
AGTTCTTCACCGAGACGGCCCATCAACACACCGGGGGATACGGAGGACTCCACGAAAGGAGGAAGCAACG
TCGCATCCGGACTACGTTCACAAGCTCCCAGTTAAAAGAGCTTGAAAAAGTTTTCGCCGAAACTCATTAC
CCGGATATTTATACAAGAGAAGAACTTGCGCTAAAAATTGATCTCACTGAAGCCAGAGTGCAGGTTTGGT
TTCAAAATCGTCGAGCAAAATGGCGAAAAATGGAGCGAGCAAAACAACAACCTCAACCCATACATTCTCC
TGGGAGTTCCCCTTCTTCTCCCAACAATATCTCGTCCATAAACAACTCGGAAAATATGATAAGTTCCGCA
AGTCCTATGGAAGATATAGGTAAGAGCAAGAGATGTATAGTT

>Phox2 C-terminus gene model (KH.C14.119)
GGTTTGCTATCTGATGACGTCACCACGCATCAATCCCATCATAATACGTCAGAAATGACGTCACCGCCCG
GTTTTCGCGGTAAACCAATCCAATCTTTGATGTTACCAAACCATGGACAATCAGGATCAAGTTATAACAG
CAATCAACAGTTCATGCAAGGTCACAACCGATCACAGCACAACACTTCATCATCGGTGTCTTGGTTCCTA
TCCGGTGGGGGTAACAACGTTGCCACCCCGCTAAATACGACTCCTTTCGCCGATGTCCTATCCGTTCTAA
CGCGTAACACACCCGGGTGCTCTACTGTCGCCGCTGCGGCAGTTTTGATGACGTCATCATCCCCCGTGAC
GTCAACTACTACGTCATACCAAAGCAGCGTAATGAGAATGAATCCACTGCCGGGGCCTAACGGTGGATTC
GACAAATCGTTCACCAACGGAGTCGGGAACAAACAGAGCAGAAATATGTTTCTGGATCAGcCCTATAGTG
AGTCGTATTA

>Pax2/5/8.a (coding sequence)
ATGAACTGGGGATCAGCAATGGCGGTTGGACCATCCAGTGTGGGACATCCATTCGTGGGATCAGGAATGG
CGGCCTCACTCACCCCTTCAAGATCAGGGCACGGTGGGGTGAACCAGTTGGGCGGGGTTTACGTGAACGG
CCGACCATTACCCGACCAAGTCCGACAACAAATAGTGGACCAAGCACACATAGGGGTTCGACCTTGTGAC
ATTGCTAGACAACTCCGGGTGTCGCATGGTTGTGTAAGCAAGATATTAGCAAGATATTACGAGACAGGCA
GCATCCGACCTGGTGTTATCGGTGGAAGCAAACCTAAGGTAGCTACACCTCGGGTAGTGGAGAAGATATG
TGATTATAAACGACAGAATCCAACTATGTTTGCTTGGGAGATACGAGACCGATTGTTGAGTGAGGGAATT
TGTGACCATGATAATGTACCCAGTGTTAGTTCGATCAATAGGATTGTCCGAAACAAAGCTGCAGAAAATG
CAAAGTCGCACCAACAACTCATGGTCCCAATGACGCCGTCATCGCTGGGTATCCACACAAACGGTCCGAT
ATTAATGGAACACGAGATTCGCGGAACGTACACGATTAACGATATATTAAGGTTACCCCACAACCCCCTT
CCCCCACATAACCCCCCCACCACTGAAACAGACCCAACACATTATAAACAAGCCGAAAATGGGATCCATT
ATAATCATAACGACAACTACAGCGAGCACATCGGGCAGGTGTCGCGCATGAACGGTTTCAATAGATTCGC
TTCATCGACCACAGCATTACCCACAATGCAACAATCAAGTCATGTGACTTCTCAGATGAATTTCGTTCGT
AAAGAAAACAAGGGATTGGATTATTCGTACGATTGTCGCGGTTCAACTAATTCACCCAACGTAGCTACTT
ACCCAGTGTTACCTCACAATCAACCTCGAGCTAACTGTGATGTCACAATCAGCCCAATGACATCACAAAC
CAACACTGCAAACGCGACAGTTTCACCCAGCAACAGTGGGGGTTACTCTGGGTCAAGTTTTGCCCCGATC
ACAACCGCGTACGCTCCGACAGGTGAGTTTGTGGATTATGGATATCAACAATACAATCAACACTGGAAGT
TTGGCCAACAGCACCATAATGATAGCAACACTGGTAAAGTTTTGAATCTTCGAGGTAAAGAACATCCGGC
AACACTGGAGATGGTCAGCGCTCAATAA

>Crls1 (coding sequence)
ATGATCAAATGCAGTTTGTCCATTCCAAGATTCCTGCAATCTAAAACTTTGAACTCAATCAAGACATTGG
AACGATGCCAACATGTAAAAAGTAATTGGGTGAAACACAGTGTTGGATGGTGCAACAATGGACATATTAA
CATAAAAACCTACAACACACTGGGACTTGGTCATCCTACTCTAAACAAGTTTCATCGTTGCCATTTAAGT
TGTATATCATCTCAGCAATGCAGATCATTCTCACTTAGTTGTAAGCTTCTATCTTCAAATGAAGAATGGG
AAAACCTGTTGGAAAATAAAAGCAAAAAGGAAAAAATTGAAAGTATTCCAGTGAAGAAAGAAAACATCTA
CACTGTCCCAAACCTACTTTCATTTTGTAGAATCGTGGTTTCCCCTTATCTCAGTTACCTTGTGCTTACT
GGTCAATCCCAAACAGCTTTAGTGTTGTGTGTGATTGCTGCCGTAACTGACATGATTGATGGACAAATAG
CTCGTACTTGGCCGTCACAACAAACAGCATTGGGTTCGGCACTTGATCCACTTGCTGATAAAGTTCTGGT
TGCTTTTCTTTCCCTTTCTCTCACATATGTCAACATTATACCATGGGCACTTACTGGTTTGTTCATTGGT
CGTGACGTCATCCTTATTCTCGCTGTGTTTTATCTACGTTACAAGACATGCCCACCCCCTGTAACTTGGG
AAAGATATTTTGATCCAAGTTTAGTTAATGTTAAACTCTCCCCAACCAACCTTAGCAAGGCAAACACTGC
GGCACAGTTGATTCTGTTATGGTGCTCTGTGGCTGCTCCTGTGTTTGGGTTTGTAGACCATGCTGCACTT
CGTTGTTTGTGGGCGTTCACTGCTTTTACTACAGTAGCTTCTGGTGTCAGTTATATCTTACAAAAAGACA
CAATTCGAATCATGAAAGAAAGCCCATGA

>FGF9/16/20 (coding sequence)
ATGTCTATGTTAACCAACATGTTAGGCCTCAGCAGCAATGTTCCGCAATCAGCAACGGGAACAAACTTTT
TAAGTTCGCTTTTACTCGCAGCCGTAGCCCGAGCGAGAGGCAAGGCAGTTACCGGACAATCGCCACCCGA
AAGCCCGTCGTCCGTCCTAGAAAGCTCGATGTCAAGAACCGAAGCAGCAAAAGTACAATCAGTTTTTTCA
GCCAAGCTTGGACGGGCAAGAAGAAAAAATTCCCTCTCTGTCGACTCCTCCACCCCCCAGAAATTAACAG
AAGAAACAATGCCTGTGGTACACGACCGACTCCCCGATCTCAGCCTTTGGAGCAAAATTGACGAGGAATT
AAAAGAGCAAGAGAAATCTTCCGCCAATGGACTATTGTCCTCAACCTCAGCAAGGAGTAAGAGAAACACC
GGTATGTCTGCCTACGACTACGATACCCAAGAAACTACCATGAGACGAAGAATGTTGTATTGTAAGAACG
GATTCAACCTTCAAATTTTACGAAACGGAAGAATATCTGGTACCCAAGAAAGCCACAATCAATACGCCGT
CCTTGAATTTATTTCTACTGGAATCGGAAGTCTCACCATCCGGGGTGTCGCAAGTGGTCTCTACCTGGCA
ATGAACAGCAAAGGAAGATTGTATGCTTCGGAAACATTCAATCGCGAGTGCATCTTCTACGAAACAGTAT
TGGAAAACAACCAGAACACGTTCGAATCATTCGCCCACAGAAATGGCTCAAAGAAATGCTACATCGCCCT

TGGTCGTCATGGAAGACCAAAGCAAGGATGCAGACTACACCCAAACAACCGACACAGCCAATTCTTGCCC
AGGAACATCGATACTGATAAAGTAAAAAACCTGTACTCTGGCCGACTCTACTGA

>EphrinA.b (coding sequence from Stolfi et al. 2011[13])
ATGCCTTTGTCAAAAGTTAATCGAATGATTTTAAGTTTGTTTATGTGCTATCTTCTCTTTGATGGGGTAC
GAAGCAAGCGGTTTAATTCGGGGACCATCACCCATGAGGTTATGTGGGATCCACGAGAAAACGGAGGGTT
TGCTTTCGGAAACGAGTTCAGCATTGAGGTTCATATGCGAGACTACATGAATATTCACTGCCCTCAGTAT
GATCAAGATGATAAAGCTAAGCTTAGTTTCATCATCTACAATGTCAGCGAACAATCATACAACAGTTGCT
CTCTCACTGATATGAAAACTGCGATTTTTAAGTGTGACAACCCAGTTAAAGGACGCAAGCTCACCACCAA
GTTTCAACGAAGAAGTCCAAACCCACTGGGATTCGTATATCAACCAAATAAAGATTATTTCTTTATGGCA
TTCAAAAAAGATGAACCTCAGAATTGCAAATCAGCGATGAAAATGAAAGTTCATGTTCTGCCGAAAAGAC
TGCACGAACACGACCGGAAACCAGTCACTGCAGGTACAACACGGTCGTCAACGCCAAGGACACTTTCGTC
AACCACTACTACTACAACAATAGCAACAACAACAACAACAACACCCCGAACAAGCCGGAAACCCCATATA
CCAAGACAGAGCTCGACAAGAAATACTCCGCCTACAAAGACGTTTAAACCTGGATATGTCCGGGGCGACG
GTGAAGGTGACGGCCCGGGTGGCGGGGTCGGTAGAATTACTTGCGCCACCACGTGGATGTTAGTTGCGTT
AGTGCTGACTGTTCTGTTACAAAACTGA

Nori Satoh gene collection clones used for other in situ probes:

Eph.c: R1CiGC16p22

EphrinA.d: R1CiGC01j20

**sgRNAS used in this study:**

Pax2/5/8.a.3.75
**gGTAGCTACACCTCGGGTAG (G+N19)**

Pax2/5/8.a.3.54
**gATCGGTGGAAGCAAACCTA (G+N19)**

Phox2.3.43
**gACGGCCCATCAACACACCG (G+N19)**

Phox2.3.69
**gCGGAGGACTCCACGAAAGG (G+N19)**

Gli.2.11
**gGTGGTAGTTCAGACTCAGG (G+N19)**

Gli.2.81
**gTTACGTCATCAGTCGACCC (G+N19)**

Control (from Stolfi et al. 2014[6])
**gCTTTGCTACGATCTACATT (G+N19)**

PCR primers for amplicon sequencing by NGS

| Gene + exon | Forward primer | Reverse primer |
|---|---|---|
| Pax2/5/8.a exon 3 | TTACGAGACAGGCAGCAT | CACAAAACACAATTTCGGGTAATT |
| Phox2 exon 3 | ATTGCACCTGGGCTAGAAGA | TATTGGTAATATTGTATGTCGCAAC |
| Gli exon 2 | GTGGAACGTCCATGTGGTC | AACATGCGAATATGTCCTATCA |

**Electroporation mixes for perturbation experiments (per 700 µL of total cuvette volume)**

Pax2/5/8.a driving GFP to look at Neck Cell Lineage

80 µg Pax2/5/8.a [-1961 to -1]>GFP

Pax2/5/8.a driving H2B:mCh & CiPhox2 Driving GFP to look at Neck Cell Lineage

20 µg Pax2/5/8.a [-1961 to -1]>H2B::mCh
80 µg CiPhox2 [-2951 to -1]>GFP

Pax2/5/8.a driving mCh & CiPhox2 Driving GFP to look at Neck Cell Lineage

80 µg Pax2/5/8.a [-1961 to -1]>mCh
80 µg CiPhox2 [-2951 to -1]>GFP

CrPhox2 Driving GFP to look at juvenile motor neurons after CRISPR/Cas9 gene editing

80 µg U6>F+EsgRNA Control or
40 µg U6>Pax2/5/8.a.2.54
40 µg U6>Pax2/5/8.a.2.75 or
40 µg U6>Phox2.3.43
40 µg U6>Phox2.3.69
40 µg Sox1/2/3>Cas9:Geminin
80 µg CrPhox2 [-2416 to +15]>GFP

CrPhox2 Driving GFP to look at cell fates after Pax2/5/8.a (tv1) overexpression

80 µg CrPhox2 [-2416 to +15]>GFP
80 µg Pax2/5/8.a [-1961 to -1]>mCh
80 µg Nut [-1155 to -1]>LacZ or
80 µg Nut [-1155 to -1]>Pax2/5/8.a (tv1)

Pax2/5/8.a variant overexpression for bulk RNA sequencing.

80 µg Nut [-1155 to -1]>LacZ or
80 µg Nut [-1155 to -1]>Pax2/5/8.a (tv1)
80 µg Nut [-1155 to -1]>Pax2/5/8.a (tv2)

Vanabin4 gene target validation and CRISPR

80 µg U6>Control F+E sgRNA or
40 µg U6>Pax2/5/8.a.2.54
40 µg U6>Pax2/5/8.a.2.75 or
80 µg Nut [-1155 to -1]>Pax2/5/8.a (tv1) or
80 µg Nut [-1155 to -1]>LacZ or
40 µg Sox1/2/3>Cas9:Geminin
20 µg Nut [-1155 to -1]>H2B:mCh

Gli expression after Pax2/5/8.a CRISPR
80 µg U6>Control F+E sgRNA or
40 µg U6>Pax2/5/8.a.2.54
40 µg U6>Pax2/5/8.a.2.75 or
40 µg Sox1/2/3>Cas9:Geminin
20 µg Sox1/2/3>H2B:mCh
80 µg Gli [-3339 to +57]>GFP

Neck specific expression of FGF and Ephrin signaling proteins
80 µg Pax2/5/8.a [-1961 to -1]>LacZ or
80 µg Pax2/5/8.a [-1961 to -1]>dnFGR or
80 µg Pax2/5/8.a [-1961 to -1]>dnEph.c or
80 µg Pax2/5/8.a [-1961 to -1]>caMEK
80 µg CrPhox2 [-2416 to +15]>GFP
20 µg Pax2/5/8.a [-1961 to -1]>H2B::mCh
80 µg Pax2/5/8.a [-1961 to -1]>mCh

Neck Specific Expression of BMP receptors
80 µg Pax2/5/8.a [-1961 to -1]>LacZ or
80 µg Pax2/5/8.a [-1961 to -1]>caBMPR or
80 µg Pax2/5/8.a [-1961 to -1]>dnBMPR
20 µg Pax2/5/8.a [-1961 to -1]>H2B::mCh
80 µg Pax2/5/8.a [-1961 to -1]>GFP

Neck morphology after Gli CRISPR
80 µg U6>Control F+E sgRNA or
40 µg U6>Gli.2.11
40 µg U6>Gli.2.81
40 µg Sox1/2/3>Cas9:Geminin
80 µg Sox1/2/3>GFP
80 µg Pax2/5/8.a [-1961 to -1]>mCh

Precocious CMNs in juvenile after Neck-specific manipulations of FGF and Ephrin signaling
80 µg Pax2/5/8.a [-1961 to -1]>LacZ or

80 μg Pax2/5/8.a [-1961 to -1]>dnFGR or
80 μg Pax2/5/8.a [-1961 to -1]>dnEph.c or
80 μg Pax2/5/8.a [-1961 to -1]>caMEK
20 μg Pax2/5/8.a [-1961 to -1]>H2B::mCh
80 μg Pax2/5/8.a [-1961 to -1]>GFP

| **Gene** | **KH ID** | **KY21 ID** | **ANISEED ID** |
|---|---|---|---|
| *Pax2/5/8.a* | S1363.2 | Chr6.690 | Cirobu.g00013823 |
| *Phox2 (Nter)* | C14.100 | Chr14.160 | Cirobu.g00003507 |
| *Crls1* | C11.724 | Chr11.285 | Cirobu.g00002581 |
| *Phox2 (Cter)* | C14.119 | Chr14.159 | Cirobu.g00003527 |
| *Gli* | C7.334 | Chr7.445 | Cirobu.g00008171 |
| *Vanabin4* | C3.88 | Chr3.256 | Cirobu.g00006092 |
| *FGF9/16/20* | C2.125 | Chr2.824 | Cirobu.g00004295 |
| *Eph.c* | C7.568 | Chr7.844 | Cirobu.g00008427 |
| *EphrinA.b* | C3.202 | Chr3.875 | Cirobu.g00005364 |
| *EphrinA.d* | C3.716 | Chr3.881 | Cirobu.g00005918 |
